# Designed NGF mimetics with reduced nociceptive signatures in neurons

**DOI:** 10.1101/2025.04.14.648806

**Authors:** Thomas Schlichthaerle, Aerin Yang, Damien Detraux, David E. Johnson, Chloe J. Peach, Natasha I. Edman, Catherine Sniezek, Charles A. Williams, Sonali Arora, Neerja Katiyar, Irene Chen, Ali Etemadi, Andrew Favor, David Lee, Connor Kubo, Brian Coventry, Buwei Huang, Stacey Gerben, Nathan Ennist, Lukas Milles, Banumathi Sankaran, Alex Kang, Hannah Nguyen, Asim K Bera, Babak Negahdari, Nobuhiko Hamazaki, Devin K. Schweppe, Lance Stewart, Jessica E. Young, Nigel W. Bunnett, Hannele Ruohola-Baker, Julie Mathieu, Siobhan S. Pattwell, K. Christopher Garcia, David Baker

## Abstract

The clinical use of Nerve Growth Factor (NGF) for neuronal regeneration has been hampered by pain sensitization side effects. NGF signals through the receptor tyrosine kinase TrkA and the co-receptor p75^NTR^; pain sensitization is thought to involve p75^NTR^. We sought to overcome this limitation by de novo design of a TrkA agonist that does not bind p75^NTR^. We designed homodimeric TrkA engaging constructs that dimerize TrkA subunits in a variety of geometries, and identified those eliciting the strongest signaling. The resulting designed agonists are able to stimulate transdifferentiated neurons and neuroblastoma cell lines, leading to neurite outgrowth and neuronal differentiation, with considerably reduced transcription of inflammation and pain related genes. These agonists are promising candidates for promoting neuronal regeneration without adverse side effects.

**Highlights:**

- De novo designed TrkA agonists activate MAPK and PI3K-AKT signaling
- Rigid fusions allow for highly tunable signaling signatures
- TrkA agonists lead to neurite outgrowth in neuroblastoma cells comparable to retinoic acid
- Modulation of the TrkA pathway without co-stimulating p75^NTR^ leads to a downregulation of inflammatory and nociceptive signature in neurons.

**Graphical Abstract:** 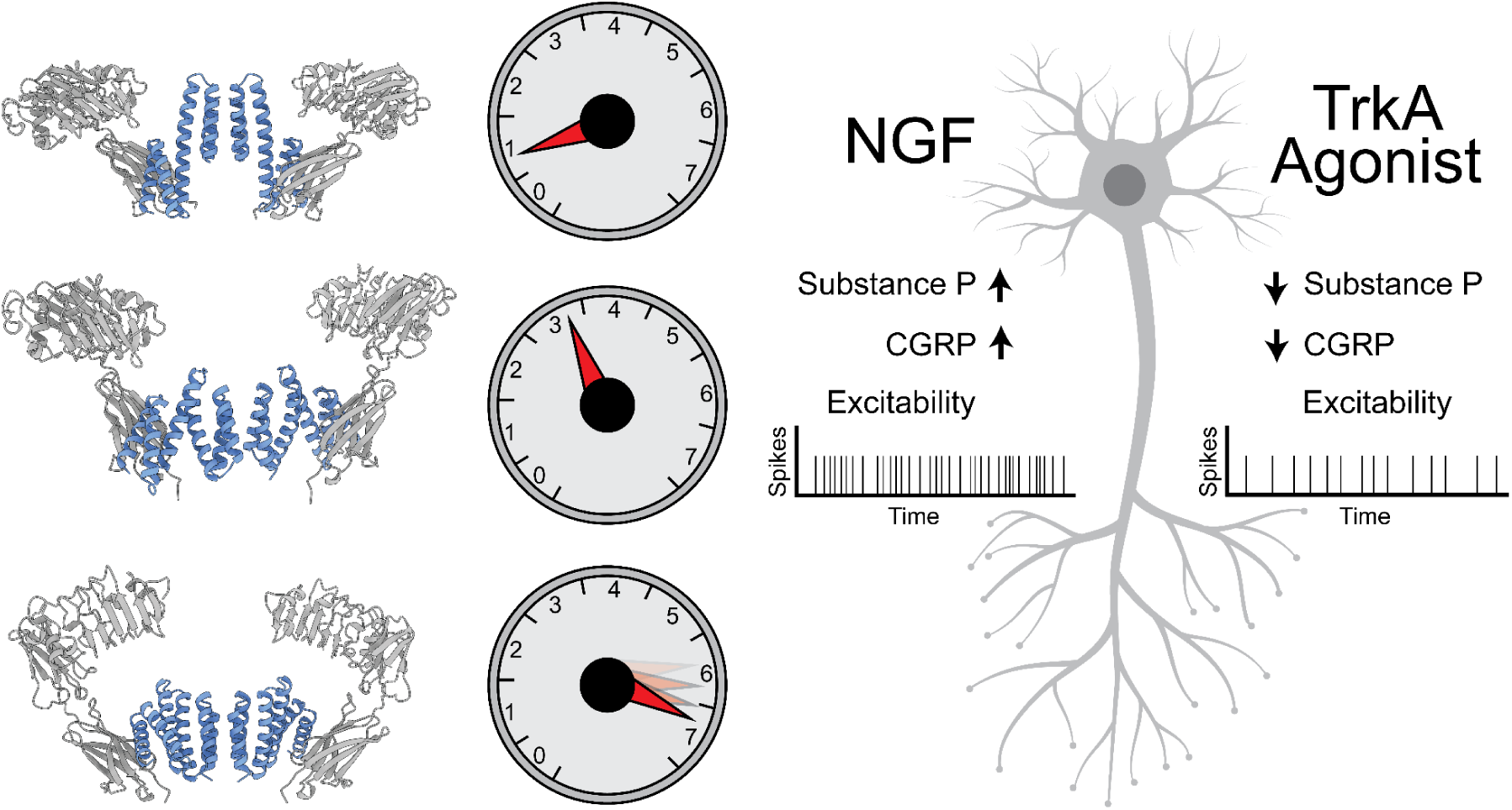

## Introduction

Nerve Growth Factor (NGF) belongs to the neurotrophin family (BDNF, NT-3, NT-4, and NGF) which play critical roles in the maintenance and development of the mammalian nervous system^1^. All neurotrophins bind to p75^NTR^, a receptor of the tumor necrosis factor family (TNFR) and preferentially engage one or more of the three tropomyosin-related receptor kinases (TrkA, -B, -C)^2^. NGF binding induces TrkA dimerization (**Figure 1A**)^3,4^, which is a common feature in receptor tyrosine kinase (RTK) signaling, leading to phosphorylation of intracellular tyrosine residues. This triggers adaptor protein recruitment and activates downstream signaling cascades such as ERK, AKT and PLCγ which plays an essential role in the modulation of sensory neurons in adults^5^. In addition to stimulating neuronal proliferation, regeneration, and differentiation, NGF induced pathway activation is also a potent mediator of inflammation and nociception upon tissue injury^6^. Notably, antibody-mediated sequestration of NGF has been shown to reduce both acute and chronic neuronal hypersensitivity in various models of inflammatory injury^7^, such as colitis^8^, arthritis^9^ or femur fracture^10^. Human genetics studies suggest this pain response is due to binding of NGF to p75^NTR^: HSAN V (human sensory and autonomic neuropathy type V) patients show a loss of pain perception^11,12^ which is mediated by a mutation in the NGF gene (R100W), which leads to decreased binding of NGF to p75^NTR^ (**Figure 1B**)^13,14^.

**Figure 1:**
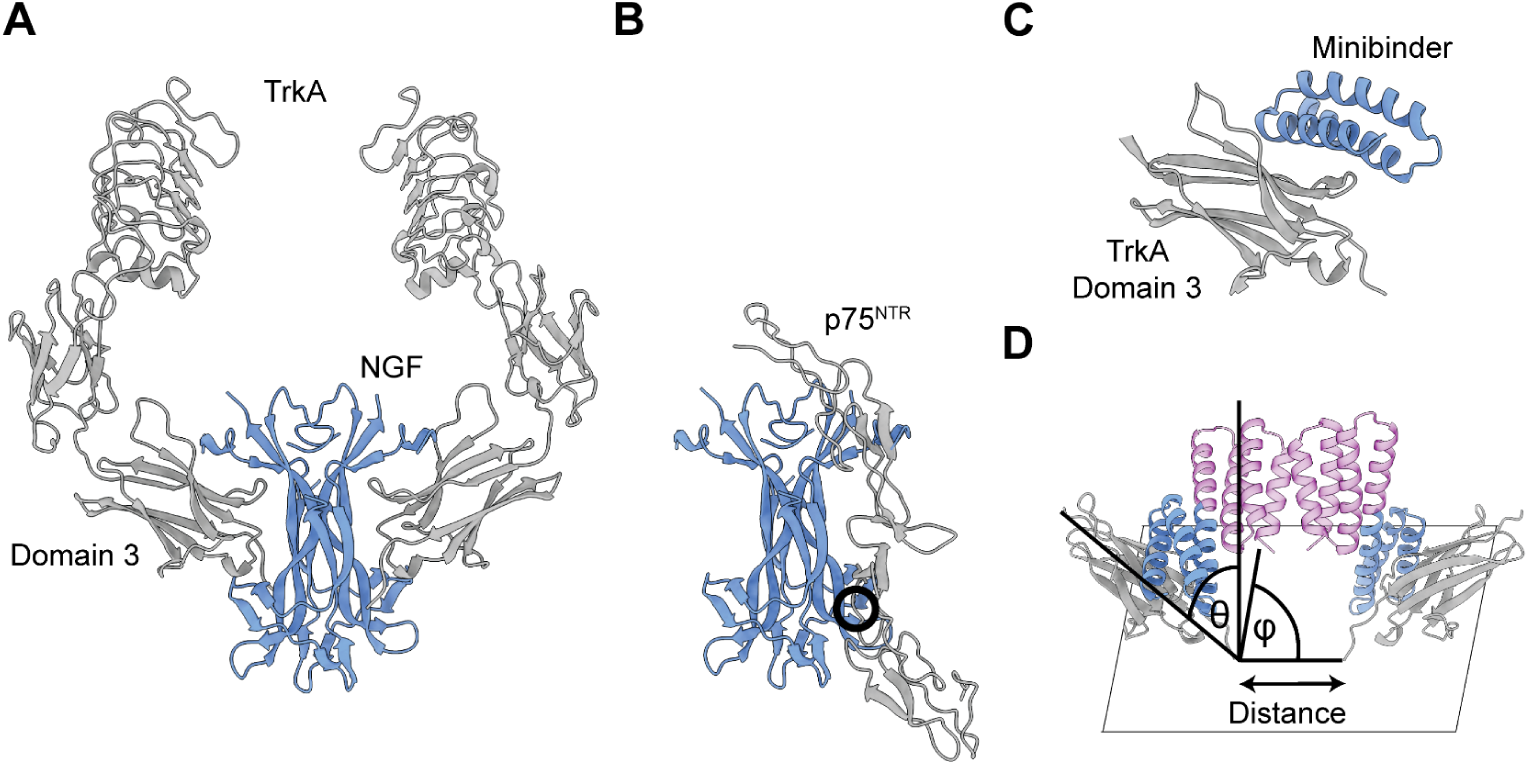
De novo design of neotrophins. (A) Native NGF-TrkA receptor complex (PDB ID: 2IFG). (B) Native NGF-p75^NTR^ complex (PDB ID: 1SG1). Black circle indicates the position of the R100W mutation which abolishes the binding of NGF to p75^NTR^. (C) De novo designed minibinder in complex with TrkA (PDB ID: 7N3T). (D) Positioning of domain 3 of the TrkA receptor via de novo designed neotrophins and exploration of three degrees of freedom: the distances between the membrane proximal receptor termini, theta (ϴ - angle between the plane of the membrane and the z-axis) and Phi (φ - angle in the plane of the membrane) were sampled.

Based on these findings, we reasoned that TrkA specific NGF mimetics (which we call Neotrophins) that do not engage p75^NTR^ could stimulate neuronal regeneration without eliciting inflammation or sensitizing pain perception. To achieve this goal, we set out to design homodimeric TrkA candidate agonists in which each subunit has an intact TrkA binding site (**Figure 1C**) that bring together two TrkA subunits to promote cross-phosphorylation of the receptor intracellular kinase domains and activation of signaling^15^.

## Results

### De novo design of receptor templating by rigid fusions

To generate TrkA specific agonists, we designed rigid homodimers in which each subunit contains a high affinity TrkA binding domain^16^ (**Figure 1C**); the two TrkA binding sites are positioned to bind two TrkA molecules in a geometry suitable for signaling. We extended the C-terminal helix of the designed binding domain, and fused it to designed homodimers either directly or via rigid designed helical repeat (DHR) spacers^17^ (**Supplementary Figure 1**)^18,19^. As part of the design campaign, we solved the X-ray crystal structures for two of the newly generated C2-oligomeric building blocks which align well with the design models (**Supplementary Figure 2**). The designed TrkA binding homodimers were modeled in complex with two TrkA subunits, and those with physically implausible geometries (clashes between the TrkA molecules, or penetration into the membrane) or geometries unlikely to signal (angle between the TrkA subunits greater than 85 degrees) were removed. This protocol yielded a diverse set of structures that position the extracellular domain 3 of two TrkA receptors in a range of distances and angles (**Figure 1D**, **Supplementary Figure 3**). Sequences were optimized for designs which passed the geometric filters using ProteinMPNN^20^, while retaining key residues in the binding interface to the target receptor and within the C2-oligomeric interface, and designs with Alphafold2^21^ pLDDT above 85 and RMSD below 2 Å to the design model were selected for experimental characterization.

### Experimental characterization

Genes encoding 170 designs were ordered and expressed in 96-well plates and purified using nickel NTA chromatography. From those, 118 designs expressed soluble and 67 ran as monodisperse peaks on size-exclusion chromatography (SEC) (**Figure 2A-D**, **Supplementary Figure 4-6**, **1st/2nd column)**. 28 designs had CD spectra matching the designed secondary structure and were stable up to 95^∘^ C (**Figure 2A-D**, **Supplementary Figure 4-6**, **3rd column**). Binding to the TrkA receptor was tested with surface plasmon resonance (SPR) and revealed K_D_s between 0.9 and 29.8 nM for the constructs, similar to the constituent TrkA binding domain alone^16^ (1.4 nM) **(Figure 2A-D, Supplementary Figure 4-6, 4th column**). Mass-Photometry^22^ was used to evaluate the assembly state of the designs (including a GFP-based mass-tag) and showed a monodisperse peak at the correct dimeric mass (**Supplementary Figure 7**).

**Figure 2:**
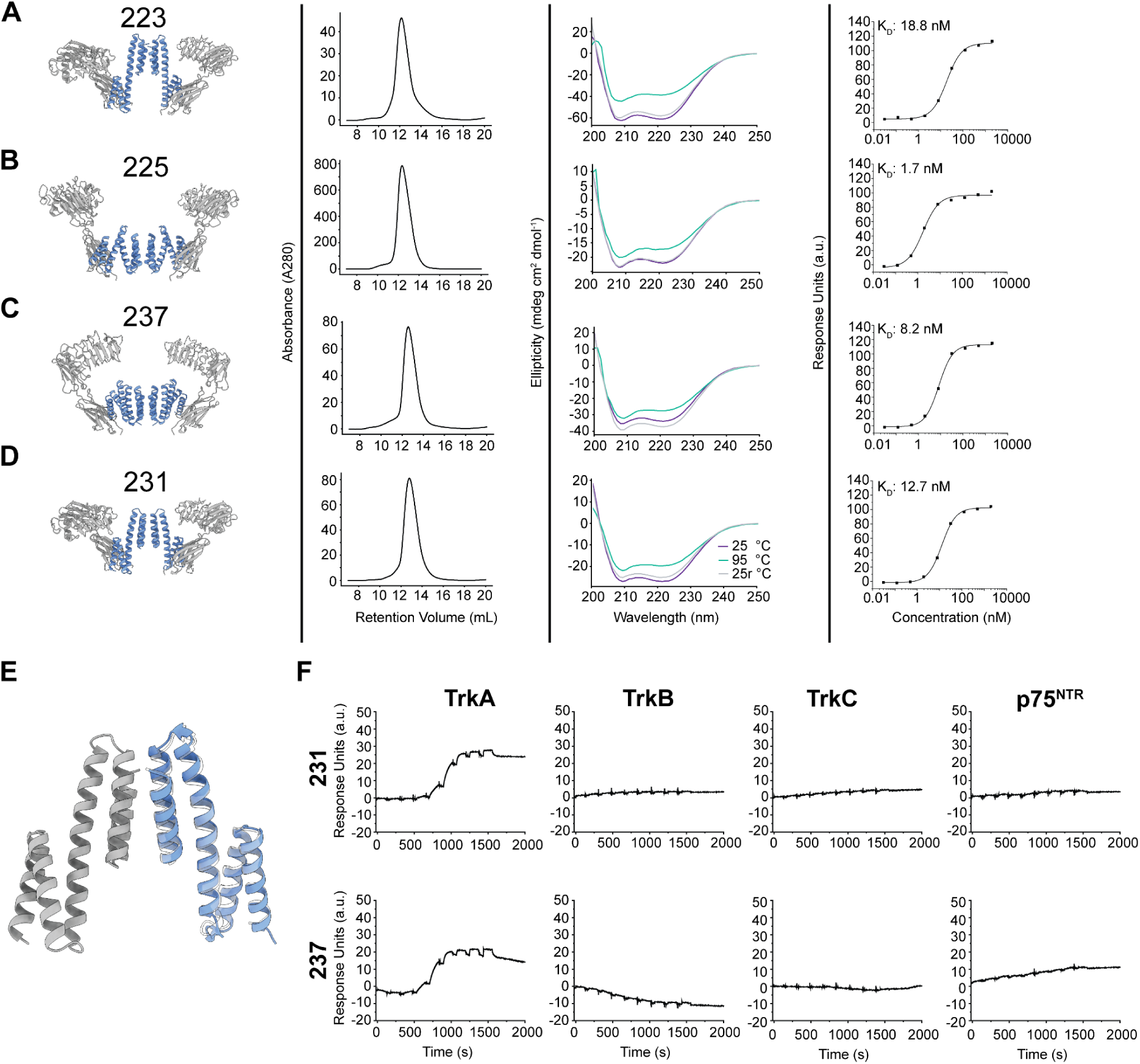
Biophysical characterization of designed TrkA receptor agonists. From left to right: cartoon model of the designed dimers superimposed with the TrkA receptor structure (PDB ID: 2IFG, 7N3T), size-exclusion chromatogram, CD spectra and dose-response curve measured by SPR for constructs (A) Nt-223 (B) Nt-225 (C) Nt-237 (D) Nt-231. (E) Crystal structure of Nt-231 constructs (blue) aligns well with the design model (grey) (PDB ID: 9DT9). (F) SPR based measurement of binding specificity of design Nt-231 and Nt-237 against TrkA, TrkB, TrkC and p75^NTR^ receptors.

We obtained a crystal structure for one of the designed constructs which directly fuses the TrkA minibinder to a C2-oligomer and positions the receptors at 39 Å, close to the native NGF mediated distance of 34 Å. The structure crystallized as a monomer rather than the dimer; the monomer aligns well with the design model with a C_ɑ_ RMSD of < 1 Å (**Figure 2E**, **Supplementary Table 1**).

To test the specificity of the neotrophins (Nt), we evaluated binding for TrkA, TrkB, TrkC and p75^NTR^. Binding was only observed for TrkA (**Figure 2F**), indicating that the designs are highly specific.

### Designed neotrophins activate signaling pathways

Designs that were well expressed, stable, and bound TrkA were tested for signaling and MAPK and PI3K-AKT pathway activation on TF-1 cells, a human erythroleukemia cell line that expresses TrkA receptors. We observed a wide range of behaviors for different design geometries, as illustrated in **Figure 3A-F** **(Supplementary Table 2)**; in these panels a representative design for each class is shown on the left, and dose response phospho-ERK and phospho-AKT curves on the right. Designs with rigid linkers (Nt-11/Nt-107) greater than 100 residues (**Figure 3A**) showed strong ERK phosphorylation (E_max_ in % NGF: 96.7 and 81.7) but had only moderate AKT activity (E_max_ in % NGF: 47.6 and 48.4), showing biased agonism towards the MAPK pathway. Six helical bundle like geometries (**Figure 3C** **and 3D**) ranged from moderate pERK activation (E_max_ in % NGF: 50.7) to strong activation (E_max_ in % NGF: 105.4), with the best agonist neotrophin 237 (Nt-237) achieving an EC_50_ of 8.7 nM for pERK. The most active designs were direct fusions with the TrkA minibinder directly fused to a C2-oligomer without helical repeat protein spacers (**Figure 3E**). Overall, the rigid fusions span a large diversity of signaling outcomes, ranging from low nanomolar EC_50s_ for pERK and pAKT pathway activation and E_max_ values rivaling NGF, down to 41.5 % E_max_ for pERK activity for the weakest design (Nt-251; **Supplementary Figure 8**). Full dose-response curves for individual neotrophins for pERK and pAKT activation can be found in **Supplementary Figure 9 and 10**, respectively. The extent of pERK and pAKT activation was reasonably correlated over the full set of designs (**Figure 3G**). We found further that most of the designs increased cell proliferation in a concentration-dependent manner (**Supplementary Figure 11**).

**Figure 3:**
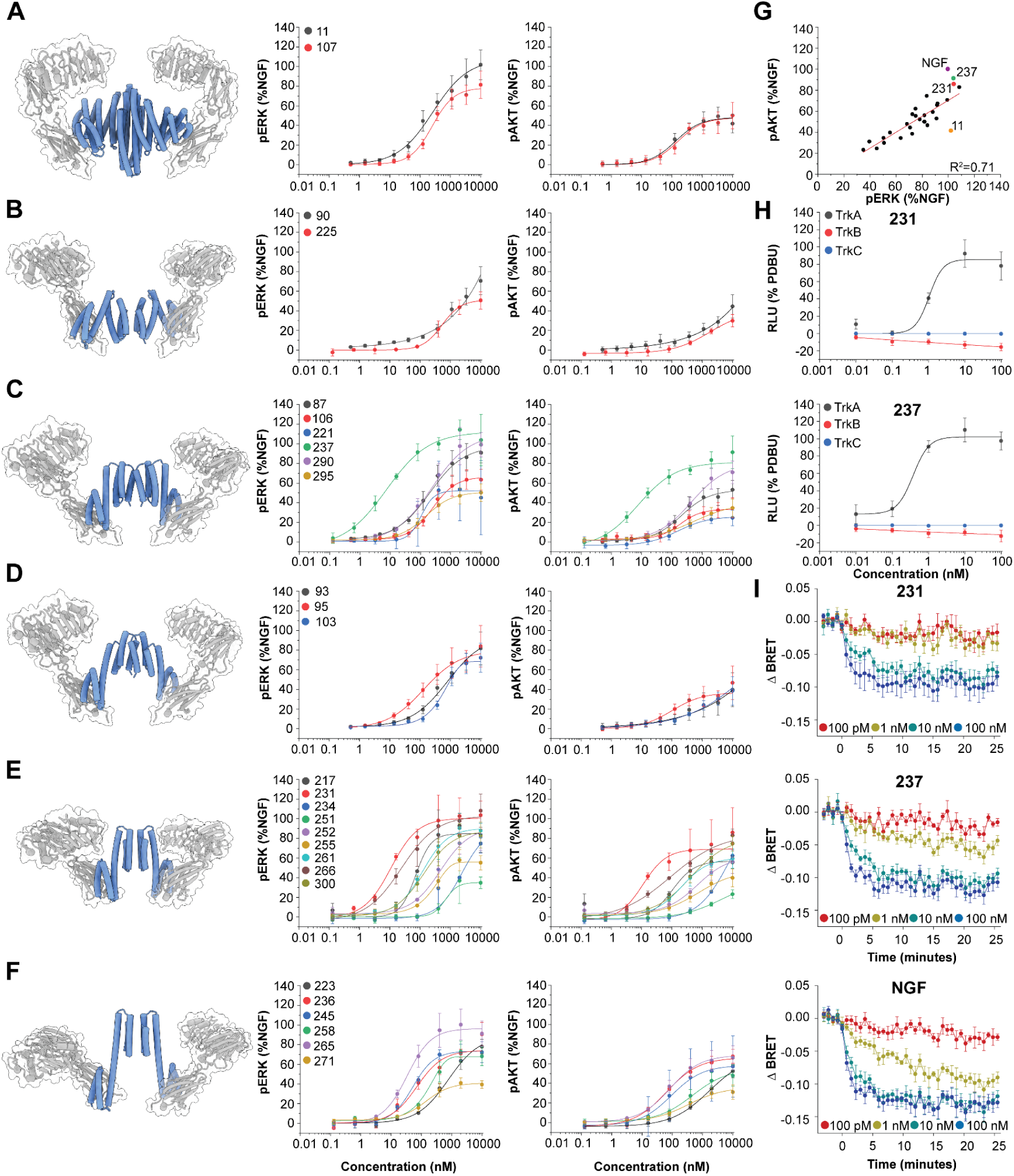
Signaling of de novo designed agonists. Dose-response curves for pERK (middle) and pAKT (right) activation measured via phosphoflow for different backbone classes of all designs (colored) (A) Large designs (B) DHR04-based (C) 6-helical bundles (D) 6-helical short connector bundles (E) direct fusions and (F) direct long fusions. Error bars represent SD from three independent biological repeats. (G) Activation strength for the individual designs correlate for ERK and AKT phosphorylation. (H) Designs Nt-231 (top) and Nt-237 (bottom) were tested for specific TrkA pathway activation using a SRE-luciferase reporter gene and treatment with different concentrations of agonists. Designs did not lead to pathway activation of TrkB or TrkC. Error bars represent SEM from five independent biological repeats. (I) Time-course of TrkA receptor endocytosis for various concentrations of Nt-231 (top), Nt-237 (middle) agonists or NGF (bottom) using a BRET assay. Error bars represent SEM from five independent biological repeats.

We moved forward with the two best performing neotrophins (Nt-231 and Nt-237) for all downstream assays. Signaling specificity was tested using a construct with a serum-response element (SRE) coupled to expression of luciferase as a transcriptional reporter downstream of ERK^23^. Nt-231 and Nt-237 induced luciferase expression (EC_50_ of 1.0 nM and 0.4 nM respectively) on TrkA-cells, but not TrkB- and TrkC-expressing cells (**Figure 3H**) showing high specificity for signalling through the TrkA receptor. RTK ligands can induce rapid receptor internalization, therefore we tested if our TrkA receptor agonists would have the same effect using luciferase-tagged TrkA receptor and a RGFP construct targeted to the plasma membrane; the BRET signal (**Figure 3I**) between the luciferase and the RGFP indicates the amount of receptor at the plasma membrane. We found that Nt-231 and Nt-237 led to a rapid decrease in BRET signal consistent with induction of endocytosis with an EC_50_ of 4.3 nM (Nt-231) and 2.4 nM (Nt-237), which is comparable to NGF-induced endocytosis (EC_50_ of 1.2 nM). Half-life of internalization of NGF was measured at 100 nM ligand concentration to be 34.7 sec, while the T_1/2_ of 100 nM Nt-231 or Nt-237 was 57.8 sec and 45.9 sec, respectively.

### Neurotrophic effects of TrkA agonists in neuroblastoma cell lines

Neuroblastoma (NB) is a pediatric cancer of the sympathetic nervous system, thought to originate from immature neural-crest derived cells of the sympathoadrenal lineage with arrested differentiation^24^. Numerous studies have utilized NB cell lines to investigate neuronal differentiation and found therapeutic applications for inducing differentiation via treatment with retinoic acid^25^. NGF-TrkA signaling can also induce neuronal differentiation, but a problem with using NGF is that it can engage multiple Trk receptors making it difficult to ascertain which Trk receptor can mediate differentiation in NB. We assessed the effects of our TrkA specific agonists on SH-SY5Y NB cell lines and observed increases in neurite length within 24 hours of treatment (**Figure 4A and B**); retinoic-acid and NGF were used as positive controls. To ensure that differences in quantification of neurite length are not due to differences in the amount of cell body clusters, we quantified and compared the number of cell body clusters at 0, 24, 48, and 72 hr post treatment and did not observe significant differences in the number of cell body clusters in PBS treated vs agonist treated SH-SY5Y cells (**Supplementary Figure 12A**). Additionally, these agonists do not cause significant changes in proliferation (**Supplementary Figure 12B**).

**Figure 4:**
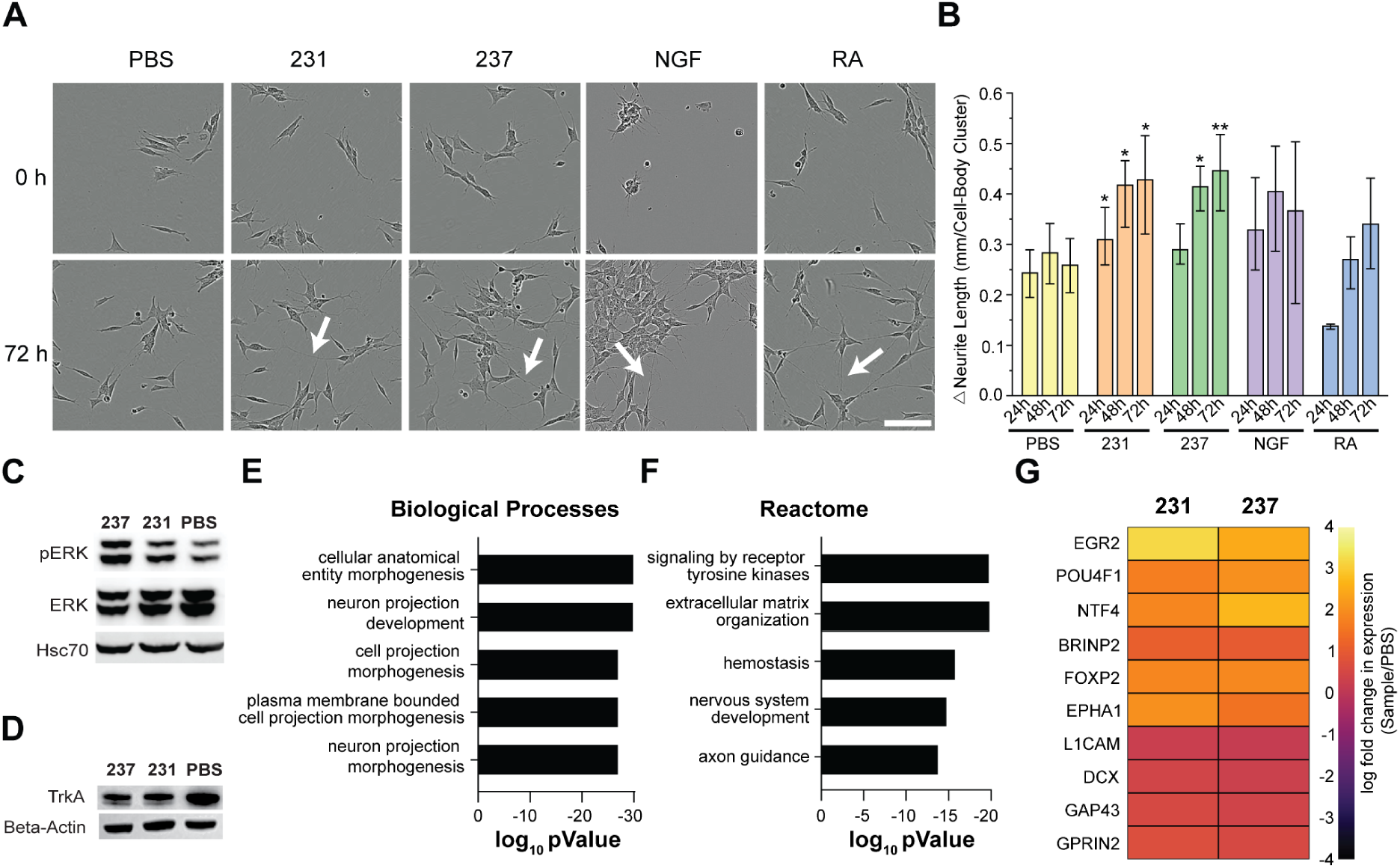
Neurotrophic effects of TrkA agonists in SH-SY5Y cells. (A) Neurite Outgrowth (white arrow) in SH-SY5Y cells treated with agonists Nt-231, Nt-237, Retinoic Acid (RA) or NGF. (B) Quantitative analysis shows that the designed agonists have strong effects on neurite outgrowth. Data normalized to the start of the experiment (Time: 0 h). Error bars represent SD from three independent biological repeats. (C) pERK signaling after stimulation. (D) TrkA downregulation after 72 hr agonist stimulation. (E) Pathway analysis after treatment of SH-SY5Y cells with agonists shows enrichment of Biological Processes relevant in neurotrophic activity. (F) Reactome analysis shows activation of RTK signaling. (G) Genes associated with neurotrophic activity are upregulated. Scale Bar: 100 µm (A)

Treatment of SH-SY5Y with neotrophin 231 and 237 for 72 hours led to an increase in pERK (**Figure 4C**). Expression of TrkA decreased over time in SH-SY5Y, IMR-32, BE(2) and LAN-1 cells after treatment, consistent with the observation that activation of Trk receptors (via binding of neurotrophins) leads to endocytosis and downregulation of the receptor (**Figure 4D**, **Supplementary Figure 12C**)^26,27^. There were no changes in expression of the neurotrophin receptor p75^NTR^, which is expected since the agonists only bind to and activate TrkA (**Supplemental Figure 12D**). Phospho-proteomics analysis showed a moderate correlation in the phosphoproteome following Nt-231 and Nt-237 or NGF treatment (**Supplementary Figure 13**).

We used RNA-seq to investigate transcriptome changes in neotrophin-treated SH-SY5Y cells. Genes up regulated in designed agonist-treated cells comprise gene ontology (GO) groups related to axon development, axonogenesis and neuron projection development, consistent with the observed neurite elongation (**Figure 4E**). Genes associated with TrkA signaling (**Figure 4F**), such as ARC and EGR2, and neuronal plasticity and differentiation markers, such as FOXP2 and POU4F1 were also upregulated (**Figure 4G**).

### TrkA agonists do not trigger an inflammatory or nociceptive signature in TrkA and p75^NTR^ expressing neurons

To assess the molecular response to neotrophin 237 in a neuronal model system we used induced neurons directly converted from fibroblasts (iNs^28^) (**Figure 5A**). First, we confirmed the expression of both TrkA and p75^NTR^ receptors in the iNs at the protein level by western blotting (**Figure 5B**; the increase of βIII-tubulin and decrease of PDGFRA expression confirm the conversion of fibroblasts into iNs). Immunofluorescence confirmed the presence of TrkA mainly on mature neurons (βIII-tubulin^+^/MAP2^+^) (**Figure 5C**). We treated the iNs with either 10 nM of Nt-237 or NGF for two days, and found that both activate the TrkA downstream pathways AKT, ERK1/2 and PLCγ1 (**Figure 5D**). RNA-seq analysis showed enrichment in biological processes related to nervous system development and the generation of neurons (**Figure 5E**), as well as Trk mediated signaling and NGF stimulated transcription patterns (**Figure 5F**) when comparing neotrophin treatment to control conditions. Comparing differentially expressed genes in iNs after NGF and neotrophin treatment showed similar upregulation of genes related to neurodevelopmental processes, such as ISL1, ROBO1 or NRP1 (**Figure 5G**) in comparison to the control condition.

**Figure 5:**
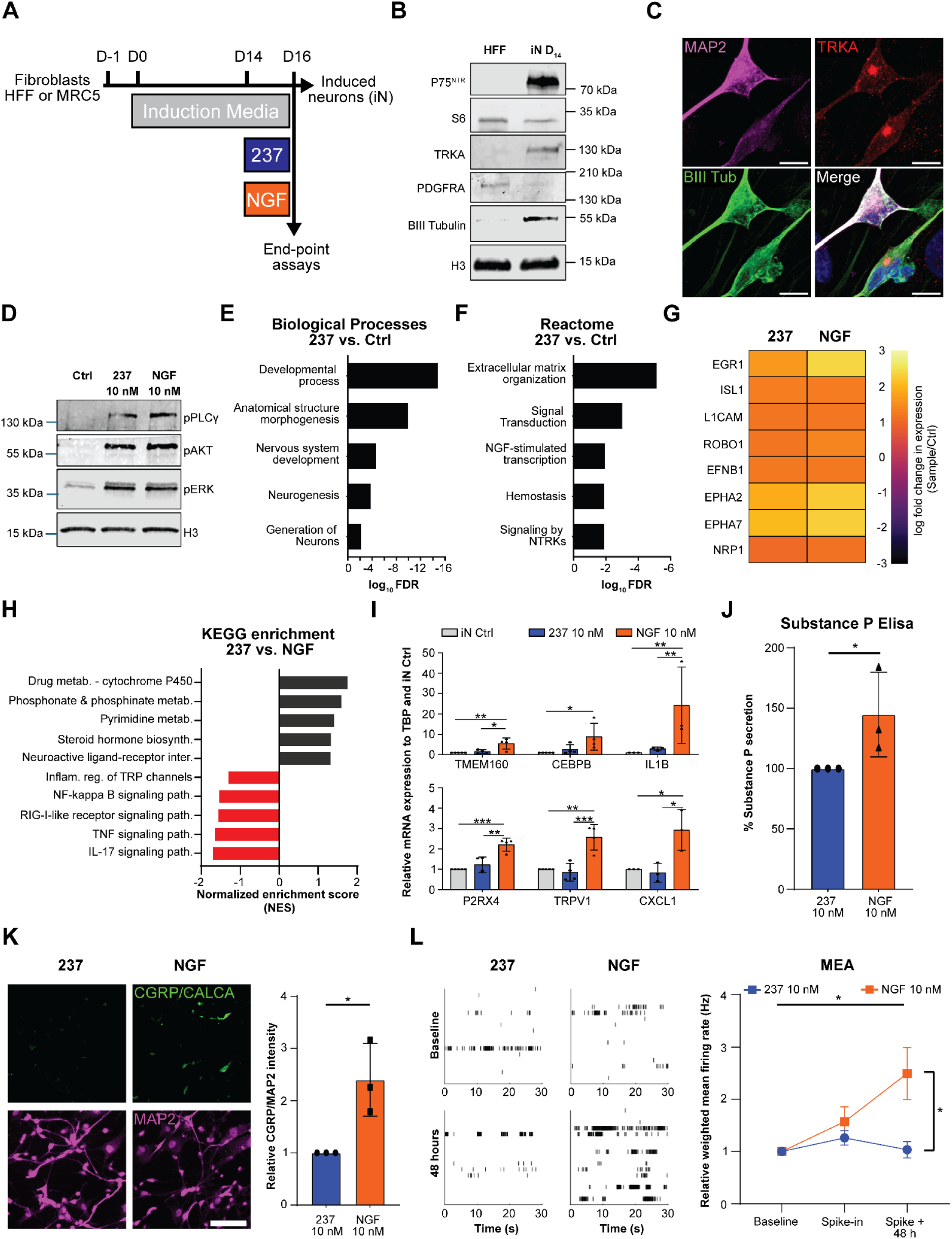
Neotrophin 237 does not trigger an inflammatory or nociceptive signature in TrkA and p75^NTR^ expressing neurons. (A) Schematic diagram depicting the timeline of fibroblast-to-neuron direct conversion and the treatment with 10 nM of either Nt-237 or NGF. (B) Immunoblot of fibroblasts before conversion and induced neurons showing expression of TrkA, p75^NTR^ and βIII-Tubulin, converted neurons show reduced PDGFRA expression. H3 or S6 are used as loading control. (C) Confocal microscopy representative image of converted neurons (MAP2+ and βIII-Tubulin+, magenta and green respectively) expressing membrane-bound TRKA (red). Dapi is used as nuclear counterstain (blue). Scale bar: 10 μm. (D) Immunoblot of D14 neurons showing activation of TrkA downstream pathways (phosphorylated ERK1/2, AKT and PLCγ) following a 15 minute treatment by 10 nM NGF or Nt-237 in starved iNs. H3 is used as loading control. (E-F) Enriched terms from RNA-seq analysis of 237-treated iNs compared to control, using Biological Processes (E) and Reactome pathways (F). (G) Heatmap of differentially expressed genes in iNs treated with Nt-237 or NGF compared to controls, shown as log fold change relative to control. (H) Top KEGG pathways enriched in Nt-237 (black) versus NGF (red) treated iNs and ordered by normalized enrichment score (NES) and with pval<0.05. (I) Validation of inflammation- (IL1B, CXCL1, CEBPB) and nociception-related (TRPV1, TMEM160, P2RX4) gene expression by quantitative real-time PCR of D16 iNs treated with either 10 nM of Nt-237 or NGF. Error bars represent SD from three independent biological repeats. (J) Quantification of Substance P release in cell culture supernatant in response to either 10 nM of Nt-237 or NGF by ELISA immunoassay. Error bars represent SD from three independent biological repeats. (K) Confocal microscopy images showing expression of CGRP (in green) in MAP2+ iNs (in purple) in Nt-237- and NGF-treated neurons. The ratio of CGRP fluorescence intensity to MAP2 intensity is quantified for normalization. Scale Bar: 100 μm. (L) Raster plots (left) and quantification (right) of electrical activity after treatment with 10 nM of Nt-237 or NGF, measured in multi electrode array plates from hiPSC-derived neurons. Error bars represent SEM from three independent biological repeats.

Since NGF activity is known to increase the expression of nociceptive-related genes (through p75^NTR^ signaling) like ASIC3^29^, TRPV1^30^ or P2X^31^ receptors, we directly compared Nt-237- and NGF-treated iNs using Gene Set Enrichment (GSEA) analysis and found specific enrichment of terms related to inflammatory and nociceptive (TRP channel modulation) responses in the NGF-treated neurons but not in the Nt-237 treated neurons (**Figure 5H**, **Supplemental Figure 14**). This is consistent with previous studies of the role of NGF in promoting inflammation^32–34^ including IL-17^35–37^ and TNF signaling^38,39^. We confirmed the inability of Nt-237 to induce the expression of genes coding for inflammatory mediators like *CXCL1, IL1B* and *CEBPB* and nociception mediators such as *P2RX4, TRPV1* and *TMEM160* by quantitative real-time PCR (**Figure 5I**)^40–42^. NGF also induces the production of secondary neuropeptides exacerbating the nociceptive signals, such as calcitonin gene-related peptide (CGRP) and Substance P, involved in the pain response^43–45^ (**Figure 5J,K**). In contrast, neotrophin 237 induced expression of neither CGRP or Substance P.

It was previously shown that NGF enhances the excitability of neurons^46^ leading to increased sensitivity to proinflammatory and nociceptive stimuli^47^. To investigate the effects of our designed neotrophins on neuronal excitability we turned to mature hiPSC-derived neurons treated with NGF or Nt-237 and analyzed the electrical activity using a multi-electrode array (MEA) plate. A significant increase in frequency of action potentials was observed two days after NGF addition indicating an increase in neuronal excitability (**Figure 5L**); this increase did not occur with the designed agonist Nt-237. Thus, our designed TrkA specific neotrophin retains the neurotrophic activity of NGF but does not trigger an increase of neuronal excitability, seen in increased pain sensitization in neurons.

## Discussion

Many secreted growth factors and cytokines, including Neurotrophins, signal through multiple shared receptors, resulting in a diverse set of pleiotropic responses. Through de novo design we generated agonists with specificity for a single receptor subunit, and by tuning the geometry of receptor engagement, we were able to finetune the strength of pathway activation. Our designed TrkA agonists signal exclusively through the TrkA receptor, mimicking the neurotrophic effects of NGF in neuroblastoma cells, but avoiding the nociceptive responses induced by NGF interactions with the p75^NTR^ co-receptor, reflected by reduced expression of Substance P and CGRP in our iN model as well as a reduced neuronal firing rate in hiPSC-derived neurons.

Recombinant NGF has shown therapeutic potential in many pathologies like Alzheimer’s disease (AD)^48–51^, traumatic brain injury (TBI)^52^, eye diseases like glaucoma^53–55^ or neurotrophic keratopathy^56,57^, or peripheral neuropathies^58,59^ but often comes with painful or inflammatory side effects. Our designs, like painless NGF^R100W^ ^48^ or chemical modulators of TrkA^60,61^, are attractive candidates for treating these pathologies without inducing pain responses. Detailed in vivo characterization will be required to confirm that the designed agonists have the regenerative activity of NGF without the pain response in vivo. The compounds described in this paper are specific for the human receptor; we are currently generating mouse TrkA agonists to explore the therapeutic utility of our approach in vivo.

More generally, given the considerable pleitropy of most native cytokines and other signaling molecules, we anticipate that designed analogues of natural signaling molecules which elicit only the subset of desired responses could be broadly useful in medicine.

## Acknowledgements

We want to thank Luki Goldschmidt and Kandise VanWormer for maintaining the computational and laboratory resources at the Institute for Protein Design. We furthermore want to thank Anka M. Pasca for exploratory experiments in brain organoids as well as Sylvia Chen for support with protein purification. We also want to thank Xinting Li for mass spectrometry support and the Cell Assay Core at the IPD for support with cell culture maintenance. We want to thank Kevin M. Jude for EM screening experiments. We also want to thank Arvind Pillai and Shingo Honda for support with Mass Photometry. This work was supported by the European Molecular Biology Organization via ALTF191-2021 (T.S.); the Advanced Research Projects Agency-Energy (ARPA-E), U.S. Department of Energy under Award Number DE-AR0001543 (GR013290: 2021), the Bill and Melinda Gates Foundation INV-043758 (GR019486, D.B.), the Defense Threat Reduction Agency Grant HDTRA1-19-1-0003 (GR008757, D.B.), the Howard Hughes Medical Institute (GR020267, D.B., K.C.G.), Spark Therapeutics (GR017125, D.B.), The Audacious Project at the Institute for Protein Design (PG117878, PG117879, PG117866, D.B.), The Nordstrom Barrier Institute for Protein Design Directors Fund (GF124659, D.B.), the Open Philanthropy Project Improving Protein Design Fund (GF129460, D.B.), The Open Philanthropy Project Universal Flu Vaccine Fund (GF129461, D.B.), the ISCRM postdoctoral fellow award (D.D) and grants from the National Institutes of Health DE033016 and DK140839 (J.M., H.R.-B), 1P01GM081619, R01GM097372, R01GM083867, NHLBI Progenitor Cell Biology Consortium (U01HL099997; UO1HL099993) SCGE COF220919 (H.R-B), and AHA 19IPLOI34760143, Brotman Baty Institute (BBI), DOD PR203328 W81XWH-21-1-0006 and Stem Cell Gift Funds (H.R-B). The National Institute of Allergy and Infectious Disease, grant R0AI160052 (GR010198, the National Institute on Aging, grant R01AG063845 (GR009173, D.B.), the Wu Tsai Protein Innovation Fund (GF151772, T.S., D.B.). The Crystallographic work is based upon research conducted at the Northeastern Collaborative Access Team beamline 24ID-C on Advanced Photon Source, which is funded by the National Institute of General Medical Sciences from the National Institutes of Health (P30 GM124165). This research used resources of the Advanced Photon Source, a U.S. Department of Energy (DOE) Office of Science User Facility operated for the DOE Office of Science by Argonne National Laboratory under Contract No. DE-AC02-06CH11357. And on National Synchrotron Light Source 2, beamline 17-ID-2. The Center for Bio-Molecular Structure (CBMS) is primarily supported by the NIH-NIGMS through a Center Core P30 Grant (P30GM133893), and by the DOE Office of Biological and Environmental Research (KP1607011). NSLS2 is a U.S. DOE Office of Science User Facility operated under Contract No. DE-SC0012704. This publication resulted from the data collected using the beamtime obtained through NECAT BAG proposal # 311950. D.E.J., acknowledges support from the Sarah Mary Hughes Foundation. This work was supported by the ALS-ENABLE grant P30 GM124169. The Advanced Light Source is a Department of Energy Office of Science User Facility under Contract No. DE-AC02-05CH11231. D.K.S. acknowledges support from the NIH/NIGMS (R35GM150919), Washington Research Foundation, the W.M. Keck Foundation, an Andy Hill CARE Distinguished Researcher Award and the Pew Charitable Trusts. This research was funded in part through the NIH/NCI Cancer Center Support Grant P30 CA015704 through a Cancer Consortium New Investigator Award to D.K.S.

## Author Contributions

T.S., N.I.E., L.S., D.B. initiated the project. K.C.G. contributed to the conceptualization and selection of agonist designs by in vitro signaling assays, T.S., N.I.E., A.E., A.F., B.C., B.H., designed and characterized proteins. A.Y. performed TF-1 cell experiments, D.D., I.C., J.M., H.R-B. performed and supervised the iN work, D.E.J., S.A., N.K., S.P. performed and supervised Neuroblastoma cell work, C.J.P. performed endocytosis and signaling specificity experiments, C.S. and D.K.S. performed and supervised mass-spectrometry experiments, C.A.W. and J.E.Y. performed and supervised MEA experiments, D.L., C.K. and N.H. performed cell experiments, S.G., L.M. purified proteins, N.E. performed CD experiments, B.S., A.K., H.N. and A.K.B. solved crystal structures, B.N., N.H., D.K.S, L.S., N.W.B., H. R-B.,J.M., K.C.G, S.P., D.B. supervised the project. T.S., A.Y., D.D., D.E.J., H.R-B., J.M., S.P., K.C.G. and D.B. wrote the manuscript with inputs from all authors.

## Declaration of interests

The authors plan to file a patent application.

D.K.S. is a consultant and/or collaborator with ThermoFisher Scientific, AI Proteins, Genentech, and Matchpoint Therapeutics.

## STAR Methods

Key resources table

Key resources table
| REAGENT or RESOURCE | SOURCE | IDENTIFIER |
| --- | --- | --- |
| **Antibodies** |  |  |
| Monoclonal Rabbit anti-phospho-p44/42 MAPK (Erk 1/2) (Thr202/Tyr204) | Cell Signaling | Cat# 4370S; RRID: AB_2315112 |
| Polyclonal rabbit Phospho-p44/42 MAPK (Erk1/2) (Thr202/Tyr204); 1:1000 | Cell Signaling | Cat# 9101S; RRID: AB_331646 |
| Monoclonal rabbit Phospho-Akt (Ser473) (D9E) XP; 1:1000 | Cell Signaling | Cat #4060S; RRID: AB_2315049 |
| Monoclonal mouse anti-p-PLC γ1 (pY783.27); 1:1000 | Santa Cruz biotechnology | Cat #sc-136186; RRID: AB_2163561 |
| Polyclonal Rabbit Anti-Histone H3 antibody; 1:5000 | Abcam | Cat #ab1791; RRID: AB_302613 |
| Polyclonal Rabbit p75 NGF Receptor / CD271 antibody; 1:1000 | GeneTex | Cat #GTX102262; RRID: AB_11162093 |
| Polyclonal chicken anti-MAP2 antibody; 1:2000 | Abcam | Cat #ab92434; RRID: AB_2138147 |
| Monoclonal Rabbit anti-TRKA recombinant (SU0354); 1:100 | ThermoFisher | Cat #MA5-32123; RRID: AB_2809414 |
| Monoclonal mouse Anti-Calcitonin Gene Related Peptide (CGRP) Antibody (4901); 1:100 | Antibodies Inc | Cat #311-CGRP; RRID: AB_2492054 |
| Monoclonal rabbit Anti-TRKA(D7U3A) | Cell Signaling | Cat #30697; RRID: AB_2798993 |
| Monoclonal rabbit Anti-p44/42 MAP Kinase | Cell Signaling | Cat #4695; RRID: AB_390779 |
| Monoclonal mouse anti-HSC70 | Santa Cruz | Cat #sc-7298; RRID: AB_627761 |
| Monoclonal mouse anti p44/42 MAPK (Erk1/2) (3A7) | Cell Signaling | Cat #9107; RRID: AB_10695739 |
| Rabbit polyclonal anti phospho-Trpv1 (Ser502); 1:1000 | ThermoFisher | PA5-64860; RRID: AB_2663797 |
| Monoclonal mouse anti-beta actin | Sigma | Cat #A1978; RRID: AB_476692 |
| Anti-mouse HRP | Azure | Cat #AC2115; RRID: AB_3661795 |
| Anti-rabbit HRP | Azure | Cat #AC2114; RRID: AB_3661796 |
| **Bacterial and virus strains** |  |  |
| E.coli BL21 (DE3) | NEB | Cat# C2527H |
| **Chemicals, peptides and recombinant proteins** |  |  |
| Fetal Bovine Serum (FBS) | Biowest | Cat# S1620 |
| B-27 supplement (50x) | Gibco | Cat #17504-044 |
| N2 supplement (100x) | Gibco | Cat #17502-048 |
| Laminin | Santa Cruz biotechnologies | Cat #sc-29012 |
| dibutryl cAMP | Santa Cruz biotechnologies | Cat #sc-201567B |
| Noggin, human recombinant | Sino Biologicals | Cat #10267-H08H |
| LDN193189 | Medchem express | Cat #HY-12071 |
| A83-01 | Medchem express | Cat #HY-10432 |
| CHIR99021 | Medchem express | Cat #HY-10182 |
| Forskolin | Medchem express | Cat #HY-15371 |
| SB-431542 | Medchem express | Cat #HY-10431 |
| FGF | R&D Systems | Cat #233-FB-025 |
| BDNF | ThermoFisher Scientific/<br>Peprotech | Cat #450-02-10UG |
| GDNF | ThermoFisher Scientific/<br>Peprotech | Cat #450-10-10UG |
| NGF protein, human, recombinant | Sino Biologicals | Cat #11050-HNAH |
| Eagle's Minimum Essential Medium | ATCC | Cat #30-2003 |
| DMEM High glucose with Glutamax | Gibco | Cat #15140-122 |
| Trypsin-EDTA | Gibco | Cat #25300054 |
| DMEM/F12 | Gibco | Cat #31330-038 |
| Neurobasal Medium | Gibco | Cat #21103-049 |
| 15% heat-inactivated Fetal Bovine Serum | Thermofisher | Cat #10082147 |
| 10% Penicillin – Streptomycin | Millipore-Sigma | Cat #P4333 |
| HBS-EP+ Buffer 10x | Cytiva | Cat #BR100669 |
| Lipofectamine 3000 | ThermoFisher Scientific | Cat #L3000001 |
| Coelenterazine purple | Nanolight | Cat #369 |
| Chemiluminescence Radiance Plus | Azure | Cat #AC2103 |
| 4% Paraformaldehyde | Electron Microscopy Sciences | Cat #157-4-100 |
| DAPI | Sigma Aldrich | Cat #10236276001 |
| Fluoromount-G Mounting Medium | ThermoFisher Scientific | Cat #00-4958-02 |
| Sera-Mag SpeedBeads - Mag4 | Cytiva | Cat #65152105050250 |
| Sera-Mag SpeedBeads - Mag6 | Cytiva | Cat #45152105050250 |
| Sep-Pak tC18 1 cc Vac Cartridge | Waters | Cat #Wat054960 |
| Fe-NTA beads | Cube | Cat #31505-Fe |
| C18 emitter column | Ionopticks | Cat #ODY3-15075C18 |
| De novo designed oligomeric proteins | This paper | Supplementary Table 3 |
| **Critical commercial assays** |  |  |
| BD Cytofix Buffer | BD Biosciences | Cat# 554655 |
| Substance P ELISA | Enzo Bioscience | Cat #ADI-900-018A |
| Aurum total RNA mini kit | Biorad | Cat #7326820 |
| iScript reverse transcription kit | Biorad | Cat #1708891 |
| RIPA Lysis and Extraction Buffer | Thermofisher | Cat #89900 |
| Pierce BCA Protein Assay Kit | Thermo Scientific | Cat #23225 |
| 10% NuPAGE Bis-Tris protein gel | Invitrogen | Cat #NP0315BOX |
| CellTiter-Glo 2.0 Cell Viability Assay | Promega | Cat #G9241 |
| TMTpro 18-plex | ThermoFisher Scientific | Cat #A52045 |
| **Deposited data** |  |  |
| Atomic Model of C2-01018 | This paper | PDB: 9DTD |
| Atomic Model of C2-02802 | This paper | PDB: 9DTE |
| Atomic Model of 231_C2_TrkA (Nt-231) | This paper | PDB: 9DT9 |
| Bulk RNA Sequencing Data | This paper | GEO: GSE292792 |
| **Experimental models: Cell lines** |  |  |
| TF-1 cells | ATCC | Cat #CRL-2003 |
| HEK293T cells | ATCC | Cat #CRL-3216 |
| SH-SY5Y cells | ATCC | Cat #CRL-2266 |
| SK-N-SH cells | Gift from Olson Lab | RRID: CVCL_0531 |
| LAN-1 cells | Gift from Olson Lab | RRID: CVCL_1827 |
| CAD cells | ECACC | Cat #08100805 |
| WTC-11 human induced pluripotent stem cells | Coriell | Cat #GM25256 |
| Human foreskin fibroblasts HFF | ATCC | Cat #SCR-1041 |
| Human lung fibroblasts MRC5 | ATCC | Cat #CCL-171 |
| **Oligonucleotides** |  |  |
| q-RT-PCR Primer Sequences | Integrated DNA | **Supplementary Table 4** |
|  | Technologies |  |
| **Recombinant DNA** |  |  |
| pET-29b(+) | Integrated DNA Technologies | https://www.addgene.org/vector-database/2569/ |
| LM627 | Addgene | Cat #191551 |
| pLVX-TREtight-Ngn2:2A:Ascl1-PGK-Puro (XTP-N2A) | Addgene | Cat #84777 |
| pLVX-EF1a-tetOn-IRES-G418 | Addgene | Cat #84776 |
| psPAX2 | Addgene | Cat #12260 |
| pMD2.G | Addgene | Cat #12259 |
| **Software and algorithms** |  |  |
| Rosetta | Koehler Leman et al.<sup>62</sup> | https://www.rosettacommons.org/software |
| Cyclic Docking Protocol | Fallas et al.<sup>63</sup> and Sheffler et al.<sup>18</sup> | https://github.com/willsheffler/rpxdock |
| ProteinMPNN | Dauparas et al.<sup>20</sup> | https://github.com/dauparas/ProteinMPNN |
| Protein Fusion Software | Hsia et al.<sup>64</sup> | https://github.com/willsheffler/worms |
| AlphaFold2 | Jumper et al.<sup>21</sup> | https://github.com/google-deepmind/alphafold |
| UCSF Chimera | Pettersen et al.<sup>65</sup> | https://www.cgl.ucsf.edu/chimera/ |
| MATLAB | Mathworks | https://www.mathworks.com/ |
| R Studio | Posit | www.posit.co |
| Origin Pro 10 | OriginLab | www.originlab.com |
| GraphPad Prism 10 | Dotmatics | www.graphpad.com |
| Fiji | Schindelin et al.<sup>66</sup> | www.imagej.net |
| Monocle3 | Trapnell et al.<sup>67</sup> | https://cole-trapnell-lab.github.io/monocle3/ |
| ComBat | Johnson et al.<sup>(81)</sup> | https://rdrr.io/bioc/sva/manual/ComBat.html |
| Adobe Illustrator | Adobe | https://www.adobe.com/products/illustrator.html |
| Axion Biosystems Neural Metric Tool | Axion Biosystems | https://www.axionbiosystems.com/ |
| STRING database | Szklarczyk et al.<sup>68</sup> | https://string-db.org/ |
| XDS | Kabsch et al.<sup>69</sup> | https://xds.mr.mpg.de/ |
| CCP4 program suite | Winn et al.<sup>70</sup> | https://www.ccp4.ac.uk/ |
| Phaser | McCoy et al.<sup>71</sup> | https://www.phaser.cimr.cam.ac.uk/index.php/Phaser_Crystallographic_Software |
| Phenix | Liebschner et al.<sup>72</sup> | https://phenix-online.org/ |
| COOT | Emsley et al.<sup>73</sup> | https://www2.mrc-lmb.cam.ac.uk/personal/pemsley/coot/ |
| MolProbity | Davis et al.<sup>74</sup> | https://www.phenix-online.org/documentation/reference/molprobity_tool.html |
| **Other** |  |  |
| SPR Biacore Series S Sensor Chip Protein A | Cytiva | Cat. #29127556 |

## Resources availability

### Lead contact

Further information and requests for resources and reagents should be directed to and will be fulfilled by the lead contact: David Baker

### Materials availability

Sequences for the generated de novo designed TrkA agonists from this study are published in Table S3.

## Data and code availability

The coordinates of the atomic models have been deposited in the Protein Data Bank under accession codes PDB: 9DTD, 9DTE and 9DT9 (C2-01018, C2-02802 and 231_C2_TrkA)
Raw and processed files for Bulk RNA-seq experiments have been deposited at GEO and are publicly available as of the date of publication. Accession numbers are listed in the key resources table.
Any additional information required to reanalyze the data reported in this paper will be shared by the lead contact upon request.

## Experimental model and study participant details

### Cell Lines

Cells were cultured under standard conditions on cell culture grade plastic dishes or flasks at 37 °C enriched with 5 % CO_2_ in incubators and generally passaged when cells reached confluency.

SH-SY5Y (ATCC, Cat. CRL-2266) were maintained in Eagle’s Minimum Essential Medium (ATCC, Cat. 30-2003) with 15% heat-inactivated Fetal Bovine Serum (Thermofisher, Cat. 10082147), and 10% Penicillin − Streptomycin (Milipore-Sigma, Cat. P4333). SK-N-SH (A generous gift from the Olson lab at Seattle Children’s Research Institute) were cultured with Eagle’s Minimum Essential Medium (ATCC, Cat. 30-2003) with 10% Fetal Bovine Serum (Thermofisher, Cat. 26-140-079), and 10% Penicillin − Streptomycin (Milipore-Sigma, Cat. P4333). LAN-1 cells (A generous gift from the Olson lab at Seattle Children’s Research Institute) were cultured with RPMI 1640 with 10% Fetal Bovine Serum (Thermofisher, Cat. 26-140-079), 10% Penicillin − Streptomycin (Milipore-Sigma, Cat. P4333), and 5% Glutamax. HEK293T (ATCC #CRL-3216) and CAD cells (ECACC #08100805) were cultured using DMEM with 10% Fetal Bovine Serum and 1% Penicillin − Streptomycin. Human foreskin fibroblasts (HFF; ATCC #SCRC-1041) and lung fibroblasts (MRC5; ATCC #CCL-171) were cultured in DMEM High glucose with Glutamax (Gibco, 10564011) supplemented with 1/100 penicillin-streptomycin (Gibco, 15140–122) and 10% FBS (Biowest)

## Method details

### Protein Design Method

#### 3 helix bundle scaffold generation

Three helical bundle scaffolds were generated by stitching together helical and turn fragments collected from previously designed miniprotein libraries. 3 helices and 2 turns were chosen at random, assembled, and checked for compactness and designability. Sequences were then assigned to the backbones using RosettaScripts and with FastDesign. The final outputs were then sub-selected using a predicted pLDDT (DeepAccNet^75^) metric > 0.9, and SecondaryStructureShapeComplementarity > 0.8.

#### Scaffold selection and cyclic docking

Subunit scaffolds consisted of a set of three helical bundles with high-resolution crystal structures. Docking was performed as previously described.^63^ Briefly, the protocol aligns subunits along the desired symmetry axis (C2) and scores these using a residue pair-motif database derived from PDB structures. Outputs were then sequence designed by Rosetta FastDesign to generate an oligomeric interface. These design outputs were filtered by ΔΔG (between -35 and -70), solvent accessible surface area (SASA > 700 A^2^), shape complementarity (sc > 0.65), and fewer than 2 unsatisfied hydrogen bonds.

#### Rigid Fusion Design

The rigid fusion designs were generated in a two-step approach. First, TrkA minibinders were fused to a previously reported library of designed helical repeats (DHRs) at the C-terminus of the minibinder. The fusion algorithm subsequently aligns a rolling window of 6 amino acids along the c-terminal helix to generate different fused structures. A filter that checks clashes to the receptor ensures reasonable designs. The library of generated fusions was then sequence designed initially with Rosetta FastDesign, later with the machine learning network MPNN. The interface residues for the previously developed minibinder to the TrkA receptor were held fixed in both design methods, to not change the binding mode and affinity. As a final step, the generated designs were predicted with AlphaFold2 (AF2). Structures that passed the AF2 filtering metrics (pLDDT > 90), were then moved to the next stage. In the second step the extended designs were fused with the same fusion algorithm (aligning and overlapping 6 amino acids of helical segments to complementary secondary structure motifs in a rolling window) to de novo designed C2 scaffolds. The structures together with the receptors symmetrized and filtered for native like geometries, including a clash-check of the newly generated fusions, a clash check of the receptors, a potential clash with the membrane (C2 designs should not be lower than the lowest point of the receptor) and native-like geometries of the receptors, which was defined as the last point in domain 3 of the TrkA receptor should not be below the first point of domain 3, to avoid receptors that would have an unnatural geometry on the cell surface. This selected for a funnel of designs (see **Supplementary Figure 1**) that is selected for native like designs. The corresponding backbones were then designed initially with Rosetta FastDesign, later with the machine learning network MPNN. The interface residues for the previously developed minibinder to the TrkA receptor were held fixed in both design methods, to not change the binding mode and affinity. Additionally, the C2 interface residues (residues to be in contact - closer then CB distance of 6 Å) were held fixed as well to not change the previously developed C2 interface. Structures were then AF2 filtered for high pLDDT (>90) and low RMSD to the design models.

#### Expression and purification

Sequences of the designed proteins were reverse translated with optimization for *Escherichia coli* expression, with a C-terminal glycine-serine linker followed by a 6x histidine tag. Sequences were ordered as synthetic genes from Integrated DNA Technologies within the pET29b+ vector between NdeI and XhoI cloning sites. This vector contains a kanamycin resistance marker and a T7 promoter. Plasmids were transformed into *E. coli* BL21 (DL3) competent cells and plated on LB with kanamycin at 50mg/L. Transformants were inoculated into 50mL of autoinduction expression media (for 1L: 12g tryptone, 24g yeast extract, 20 mL 50×M, 20 mL 50×5052, 2 mL 1M MgSO4, 200 µL Studier Trace metals, 100 µg kanamycin, q.s. to 1 L with filtered water) in a 250 mL flask. Expression cultures were grown for 20 hours at 37 °C with 200 rpm shaking. Cells were pelleted by centrifugation at 4000xg and resuspended in a lysis buffer consisting of 25mM Tris pH 8, 300mM NaCl, and 20mM imidazole with added protease inhibitor and DNase. Cells were lysed by sonication at 85% amplitude with 8 x 15 second pulses. Lysate was separated into soluble and insoluble fractions by centrifugation at 18,000xg. Immobilized metal affinity chromatography (IMAC) was used to purify designed protein. Nickel-nitrilotriacetic acid (Ni-NTA) resin was initially equilibrated with 5 column volumes (CV) lysis buffer. Supernatant was poured over the columns, followed by 20 CV wash buffer (25 mM Tris pH 8, 400 mM NaCl, 30 mM imidazole). Protein was eluted using 5 CV elution buffer (25 mM Tris pH 8, 300 mM NaCl, 500 mM imidazole). Eluate was purified by size exclusion chromatography (SEC) on an AKTA PURE FPLC system, using either a Superdex 75 Increase 10/300 GL column or a Superdex 200 Increase 10/300 GL column, with Tris-buffered saline (TBS; 25 mM Tris pH 8, 150 mM NaCl) at a speed of 0.75 mL/min. Fractions corresponding to the peak trace were collected and combined for further analysis.

#### Low-endotoxin protein production

Genes were expressed as described above. Cultures were resuspended and lysed in a phospho-buffered saline (PBS)-based lysis buffer with added protease inhibitor and DNase. Cells were sonicated and pelleted as described above. Supernatant was filtered through a 0.45µm filter prior to loading onto IMAC columns. IMAC columns were pre-washed with PBS + 1% Triton X-100 + 0.75% CHAPS to remove any residual endotoxin and equilibrated with PBS + 5 mM imidazole. Supernatant was poured onto the column and followed by washing with 5 CV PBS + 30 mM imidazole. To remove endotoxin, 4 wash steps were performed using 5 CV PBS + 1% Triton X-100 + 0.75% CHAPS, with 30 min 37 °C incubations on the first and third wash. This was followed by 2 washes with 10 CV PBS, then elution with 5 CV PBS + 400 mM imidazole. SEC was performed as described above on a dedicated AKTA PURE FPLC with lines, loops, and fraction dispenser pre-washed using 500 mM NaOH + 0.75% CHAPS. Endotoxin levels were measured with the LAL endotoxin testing system (Charles River Laboratories).

Proteins expressed by the General Protein Production core were transformed as above then a pre-culture was inoculated into 50 mL of LB media and grown at 37°C for 18 hours. 10 mL of this pre-culture was used to inoculate 500 mL of autoinduction expression media (recipe above) in a 2 L flask. Cells were lysed using a Microfluidics M-110P microfluidizer. Soluble and insoluble portions of the lysate were separated at 17000xg. Supernatant was flown over 3 mL of nickel resin and washed with PBS wash buffer (20 mM NaPO4, 300 mM NaCl, 30 mM Imidazole, 0.75% CHAPS) for 6 washes of 10mL each (a total of 60 mL). SEC was performed as described above. Endotoxin levels were measured as above.

#### High-throughput Protein Expression in 96-well plate format

High-throughput 96-well plate protein expression was performed as previously reported. In brief, Golden Gate subcloning reactions of IDT eblock based DNA strands (LM627 plasmid, including MSG-[protein]-GSGSHHWGSTHHHHHH - C-terminal SNAC tag for cleavage) were carried out in 96-well PCR plates in 5 µl volume. Reaction mixtures were then transformed into a chemically competent expression strain (BL21(DE3)) via heatshock and after a 1-hour outgrowth phase in 100 µl SOC Media was transferred into a 96-deep well plate with a total volume of 1 ml and grown overnight at 37 °C. The next day, the culture was split in 4x 96 deep well plates in a 1:10 dilution in 1 ml TB2 media supplemented with Kanamycin for a total volume of 4 ml. After 90 min at 37 °C, 0.5 mM IPTG was added to the cells and cells were grown for another 3-4 h at 37 °C with shaking. Cells were harvested and lysed with Bugbuster Master Mix. After removal of cell debris via centrifugation, clarified lysates were applied directly to a 75 µl bed of Ni-NTA agarose resin in a 96-well fritted plate and equilibrated with Tris wash buffer. The fritted plate was mounted on a vacuum manifold for faster flow through. After sample application and flow through, the resin was washed 3x with 400 µl and samples were eluted in 200 µl of Tris elution buffer containing 500 mM imidazole. Eluates were sterile filtered with a 96-well 0.22 µm filter plate prior to size exclusion chromatography.

Proteins were purified either on an Akta Pure FPLC with a 10/300 S75 column at 0.7 ml/min equipped with an autosampler and fraction collector or on an Agilent HPLC through a 5/150 S75 column at 0.6 ml/min. The run was monitored using A280 absorbance measurement, running buffer was 25 mM Tris, 100 mM NaCl at pH 8.0.

#### Surface Plasmon Resonance

Designed neotrophins were tested for binding against TrkA, TrkB, TrkC and p75 (NGFR) using surface plasmon resonance (SPR) measurements (Cytiva, Biacore 8K). In brief, 1-2 mg/ml of FC-tagged Receptor were immobilized on a Protein A chip and 9 concentrations per each design (diluted in 1xHBS-EP+ buffer) were measured with a multi-cycle kinetics setup, which titrates 9 different concentrations of binders over the chip surface. Kinetic fits and dose-response curves were measured to fit the affinity values for the individual constructs via the Cytiva Biacore software.

#### Circular Dichroism

Ultraviolet circular dichroism measurements were carried out with a JASCO-1500 instrument equipped with a temperature-controlled multi-cell holder. Wavelength scans were measured from 260 to 190 nm at 25 to 95 °C, in eight 10 °C steps. Stability was assayed by a reverse folding ramp in eight 10 °C steps from 95 to 25 °C. Wavelength scans and temperature melts were performed using 0.4 mg/ml protein in PBS buffer (20 mM NaPO4, 150 mM NaCl, pH 7.4) with a 1 mm pathlength cuvette.

#### Mass Photometry

Mass photometry measurements were carried out on a TwoMP (Refeyn) Mass photometer. To circumvent the detection limit of mass photometry, designs were ordered with N-terminal GFP tags. Before measurement samples were diluted to 1-10 nM in 25 mM Tris/150 mM NaCl and 10 ul of the diluted sample was added to one well of a 24-well gasket on the microscope slide. After autofocusing via the AcquireMP software, 1 min videos were acquired. Ratiometric contrast values for detected particles were measured and processed into mass distributions with DiscoverMP. To convert ratiometric contrast values of individual particles into mass distributions 20 nM of Beta-amylase, which consists of monomers (56 kDa), dimers (112 kDa) and tetramers (224 kDa) was used as a calibration sample. Mass distributions were plotted and fitted using a Gaussian fit with the DiscoverMP software.

#### Phospho-Flow Signaling Assay

TF-1 cells (ATCC CRL-2003) were serum-starved overnight in serum-free media without NGF or other cytokines prior to the signaling assays. The cells were then seeded into 96-well plates and stimulated with varying concentrations of human beta-NGF (R&D) or neotrophin TrkA agonists for 10 minutes at 37°C. Following stimulation, cells were fixed with 1.6% paraformaldehyde for 10 minutes at room temperature. Cells were permeabilized by resuspension in ice-cold methanol and stored at -20°C until analysis by flow cytometry. For intracellular staining, the permeabilized cells were washed and incubated with Alexa Fluor 488-conjugated anti-ERK1/2 pT202/pY204 antibody (BD) and Alexa Fluor 647-conjugated anti-Akt pS473 antibody (Cell Signaling Technology) for 1 hour at room temperature. Afterwards, the cells were washed with autoMACS running buffer (Miltenyi), and fluorescence intensity for each antibody was measured using a CytoFlex flow cytometer (Beckman Coulter). Mean fluorescence intensity (MFI) values were background subtracted and normalized to the MFI of 100 ng/ml NGF. Data were plotted in Prism 10 (GraphPad), and dose-response curves were generated using the “sigmoidal dose-response” analysis. Data were fit using a Hill function in OriginPro 2023b (10.0.5.157). Data were normalized on 100 ng/ml NGF stimulation.

#### Cell Proliferation Assay

TF-1 cells were seeded into 96-well plates and cultured in RPMI-1640 media supplemented with 2% FBS and various concentrations of TrkA agonists and NGF for 48 hours at 37°C in a humidified incubator with 5% CO_2_. Cell proliferation was assessed by measuring ATP levels using the CellTiter-Glo 2.0 Cell Viability Assay reagent (Promega) following the manufacturer’s instructions. Luminescent signals were recorded using a SpectraMax Paradigm plate reader, and the resulting data were analyzed and plotted using Prism 10 (GraphPad). Dose-response curves were generated utilizing the “sigmoidal dose-response” analysis. Data were fit using a Hill function in Origin Pro 2023b (10.0.5.157). Data were normalized on 100 ng/ml NGF stimulation.

#### Endocytosis BRET Assay

C-terminally tagged *Renilla* luciferase (Rluc8)-tagged TrkA (TrkA-HA-Rluc8) cloned with a GSSGAIA spacer using Gibson Assembly. *Renilla* Green Fluorescent Protein (RGFP)-tagged prenylation CAAX box of KRas (RGFP-CAAX) was obtained from M. Bouvier (Université de Montréal). Neuron-like CAD cells^76^ in white 96-well plates were transfected with TrkA-HA-Rluc8 (125 ng/well) and RGFP-CAAX, a fluorescent marker of the plasma membrane (150 ng/well). CAD cells were transfected with Lipofectamine 3000 (according to manufacturer’s instructions) diluted in serum-free DMEM. Following 48 h, cells were washed in Hank’s Balanced Saline Solution (HBSS) containing 0.1% Bovine Serum Albumin (BSA), then incubated with substrate coelenterazine purple (25 µg/ml, 10 min, Nanolight). Half-life of receptor internalization was calculated from the time-course datasets using a Hill fit in OriginPro 2023b (10.0.5.157) for the different constructs at 100 nM concentration.

#### SRE-Luc2P ERK Firefly Luciferase Studies

HEK293T cells plated in white 96-well plates were serum-starved following 24 and transfected with SRE-Luc2P (40 ng/well) and either NanoLuc-TrkA, NanoLuc-TrkB or NanoLuc-TrkC (20 ng/well) using PEI diluted in 150 mM NaCl solution. NanoLuc constructs had an N-terminal interleukin-6 signal sequence (IL-6) for cell surface expression. Following 48 h, cells were incubated in HBSS/0.1% BSA and stimulated with increasing agonist concentrations (10 pM-100 nM) or positive control phorbol 12,13-dibutyrate (PDBu, 10 µM). After 5 h stimulation (37°C), cells were incubated with luciferin (1:2) and lysis buffer (1:5). After 5 min to enable the substrate reaction, luminescence emissions were recorded (CLARIOstar). Concentration-response data fit using non-linear regression analysis as log *vs.* response (three parameters) to determine EC_50_ values.

#### Neuroblastoma differentiation

##### Immunoblotting

Lysates were prepared in RIPA Lysis and Extraction Buffer (Thermofisher, Cat #89900) and protein concentrations were determined by Pierce BCA Protein Assay Kit (Thermo Scientific, Cat. 23225). Proteins were run on a 10% NuPAGE Bis-Tris protein gel (Invitrogen, Cat.NP0315BOX). After blocking the membrane in 5% milk, 0.1% Tween, 10 mM Tris at pH 7.6, 100 mM NaCl, primary antibodies TrkA (Cell Signaling, Cat. 30697 at 1:1000), ERK (Cell Signaling, Cat. 9107), pERK (Cell Signaling, Cat. 4695), HSC70 (Santa Cruz, Cat. sc-7298; at 1:10.000), and Beta-Actin (Sigma, Cat. A1978 at 1:10,000) were added. Anti-mouse (Azure, Cat. AC2115) or rabbit-HRP conjugated antibodies (Azure, Cat. AC2114) were used to detect desired protein by chemiluminescence with ECL (Azure, Cat. AC2103).

##### Neurotrack

Using a flat-bottom 6-well well plate, cells were plated at 2.0 x10^5^ cells per well, in 2mL media with 100nM of either PBS, agonists 231 or agonist, 237. Using the IncuCyte^®^ Live-Cell Analysis System and IncuCyte^®^ Neurotrack Analysis Software Module enabled segmentation and quantification of neurites vs cell body clusters. Using 6-well plates, 49 images per well/condition were taken every 2 hours for 72 hrs. For analyses, 3 runs were analyzed, and an ordinary one-way ANOVA was used to determine significance.

##### Gene Ontology Analysis

Took differentially expressed genes from SH-SY5Y cells (PBS vs 231) and SH-SY5Y cells (PBS vs 237), and used Gene Ontology (https://geneontology.org/) to identify neural-related pathways.

##### Cellular Proliferation Experiment

SH-SY5Y cells were plated at 4 x10^6^ in 10 cm plates. The following day, 100nM of Nt-231, Nt-237 or PBS was added and incubated for 72 hours. After 72 hours, cells were collected and counted via a countess cell counter to assess the number of cells. This experiment was repeated in triplicate; for analysis, the replicates were averaged and the data was normalized.

##### Bulk RNAseq Analysis

Raw sequencing reads were checked for quality using FastQC^77^. RNA-seq reads were aligned to the hg38 assembly using gencode v39 and STAR^78^ and counted for gene associations against the UCSC genes database with HTSeq^79^. RSeQC^80^ were used to check for the quality of the alignments. Differential Expression analysis for RNASeq Data was performed using R/Bioconductor package DESeq2^81^ and edgeR^82^. A log2fold change cutoff of 0.32 (fold change of 25%) and FDR < 0.05 was used to find transcriptionally regulated genes in response to treatment.

##### Enrichment Analysis & Data visualization

Gene Set Enrichment Analysis (GSEA) was performed against the MsigDB database with the GO Biological Process, GO Cellular Component, GO Molecular Function, KEGG, BioCarta, and Reactome gene sets using enrichR^83^ package. Resulting enriched genesets and pathways were filtered via a threshold of FDR < 0.05. R package pheatmap (https://cran.r-project.org/web/packages/pheatmap/index.html) was used to make heatmaps. Volcano plots were made using R (v4.3.3). All other plots were made using ggplot2.

#### Cell culture and direct conversion

Human foreskin fibroblasts (HFF; ATCC #SCRC-1041) and lung fibroblasts (MRC5; ATCC #CCL-171) were cultured in DMEM High glucose with Glutamax (Gibco, Cat #10564011) supplemented with 1/100 penicillin-streptomycin (Gibco, Cat #15140–122) and 10% FBS (Biowest). Cells were passaged every 3-4 days using trypsin-EDTA (Gibco, Cat #25300054), maintained at 37°C in a humidified incubator and routinely tested for mycoplasma. For conversion to neurons, stable lines were generated by transducing two lentiviruses produced in HEK293FT cells with packaging plasmids psPAX2 and pMD2.G following the Addgene protocol: The first one containing pLVX-TREtight-Ngn2:2A:Ascl1-PGK-Puro (XTP-N2A), a gift from Fred Gage (Addgene plasmid # 84777), a the second one a tet-operator plasmid (Addgene # 84776), followed by 4 days of puromycin (1µg/ml) and G418 (150 µg/ml) selection.

The direct conversion of fibroblasts into neurons (iNs) was performed following a previously published protocol^28^. In brief, cells were plated at a density of 1×10^5/cm^2^ in regular growth media and on 0.1% gelatin-coated dishes. On day 0, cells were rinsed with PBS before switching to the neural induction media. The induction media consists of a base of a 1:1 mixture of DMEM/F12 (Gibco, 31330–038) and Neurobasal Medium (Gibco, 21103–049) supplemented with 1 x N-2 Supplement (Gibco, 17502–048), 1 x B-27 Supplement (Gibco, 17504–044) and 1/100 penicillin-streptomycin (Gibco, 15140–122). This base is supplemented with 2 µg/ml doxycycline, 1 µg/ml laminin (Santa Cruz biotechnologies), 500 µg/ml dibutyryl cAMP (Santa Cruz Biotechnologies), 150 ng/ml Noggin (SinoBiologicals), 0.5 µM LDN193189 and 0.5 µM A83-01, 3 µM CHIR99021, 5 µM forskolin and 10 µM SB-431542, all from Medchemexpress. The media was refreshed every 3 to 4 days. On day 14, 10 nM of NGF (Sinobiologicals) or Nt-237 were added. Experiments were conducted 48h later.

#### RNA isolation, qPCR and RNAseq

Total RNA was extracted directly from culture wells with the Aurum total RNA mini kit (Biorad) following the manufacturer’s protocol. Reverse transcription (RT) was performed with the iScript cDNA synthesis kit (Biorad) to convert 500 ng of RNA into cDNA.

The qPCR was performed on the ABI 7300 Real time PCR machine (Applied Biosystems) with 10 ng of cDNA per reaction, Powerup SYBR green master mix (ThermoFisher) and primers listed in **Supplementary Table 4** at a final concentration of 300 nM. Altogether, 5 ng/μl of cDNA and 300 nM of each reverse and forward primer were added per each well. Relative expressions were calculated using the 2^-ΔΔCt^ method with *TBP* as an endogenous control and untreated neurons as relative control.

250 ng of RNA were used for bulk RNA sequencing, with 20 million reads/sample, and performed by Azenta. FASTQ files were further analyzed for Gene specific analysis using *Partek™ Flow™ version 12.3.0.* For gene enrichment analysis, the StringDB online tool^68^ was used.

#### Immunocytochemistry

Cells were seeded on Matrigel-coated glass coverslips #1 (Fisher scientific) and conversion to neurons was carried out as described, before fixation with 4% paraformaldehyde (Electron Microscopy Sciences) for 15 min. Coating was performed for 1 h with 1/100 diluted Matrigel (Corning). Cells were permeabilized and blocked for 1 h in blocking buffer (PBS, 0.1% Triton X-100, 2% BSA). Immunostaining was performed by an overnight incubation of cover slips at 4 °C on 30 μL drops containing primary antibody diluted in blocking buffer. After three washes of 5 min in PBS, a second drop-based incubation was performed for 1 h in the dark and at RT with Alexa-conjugated secondary antibodies and DAPI (Sigma, 10 236 276 001) diluted 1:500 in blocking buffer. Cover slips were mounted using Fluoromount G (ThermoFisher) after three washes of 5 min in PBS. Analyses were performed with a Leica SP8 point scanning confocal (Leica microsystems) or Nikon Yokogawa W1 spinning disk confocal.

#### Western blotting

Induced neurons were starved in a pure Neurobasal media for 4 h, before being treated with 10 nM of NGF or Nt-237 for 15 minutes. The cells were rinsed in PBS before adding RIPA lysis buffer lysis buffer containing phosphatase inhibitor and Benzonase (Millipore). The surface was scraped and harvested in microtubes. The protein concentration was measured using the Biorad protein assay dye reagent following the manufacturer’s protocol (Biorad). Equal amount of proteins were mixed with 4x Laemmli buffer (Biorad), heated for 5 mins at 95°C before loading on 4-20% mini-protean gels (Biorad). The proteins were then separated for 40 mins at 200V. At the end of the migration, the proteins were transferred onto nitrocellulose membranes following the Transblot Turbo (Biorad) recommended protocol. The membranes were blocked for 1 h with a 5% BSA solution prior to an overnight incubation with the primary antibody in 5% BSA and 0.1% Tween 20, at 4°C. After three 5-minute washes in TBST, fluorescent secondary antibodies (Licor) were incubated at RT in the dark. Membranes were then washed 3 times in TBST, dried and imaged on an Odyssey M Imaging system (Licor).

#### ELISA

Concentration of Substance P in the media supernatant was obtained using a commercially available colorimetric enzyme immunoassay kit (Enzo, ADI-900-018A) following the manufacturer’s protocol. The conditioned media was harvested following 48h of treatment with either 10 nM of Nt-237 or NGF of 14 days iNs, centrifuged for 3 mins at 500g and stored at -80°C.

#### MEA

For multi-electrode array measurement, human induced pluripotent stem cells (hiPSC) derived neurons were used as previously described^84^. They were generated following the dual-SMAD inhibition^85^. In brief, hiPSCs were cultured on Matrigel coated plates and treated with neural medium (1:1 DMEM/F12 + glutamine media/neurobasal media, 0.5% N2 supplement, 1% B27 supplement, 0.5% GlutaMax, 0.5% insulin-transferrin-selenium, 0.5% NEAA, 0.2% β-mercaptoethanol) supplemented with 10 mM SB-431542 and 0.5 mM LDN-193189 (Biogems). After nine days, cells were passaged and maintained in neural medium without SMAD inhibitors. On day twelve, 20 ng/ml FGF (R&D Systems) was added to the medium and expanded. Following expansion, neural precursor cells (NPCs) were isolated using FACS, targeting positive expression of CD184/CD24 and negative expression of CD44/CD271 cell surface markers. NPCs were further expanded and differentiated into cortical neurons using neural medium supplemented with 0.02 µg/ml brain-derived neurotrophic factor (BDNF, Peprotech), 0.02 µg/ml glial-derived neurotrophic factor (GDNF, Peprotech), and 0.5 mM dbcAMP (Sigma Aldrich). Neuronal differentiation medium was changed twice weekly for three weeks. Following cortical neuron differentiation, neuronal cultures were passaged to 48 well Cytoview multiple electrode array (MEA) culture plates (Axion Biosystems) utilizing high density droplet plating. Briefly, MEA culture plates were coated with 10 µl of Matrigel positioned over the recording electrodes one day before neuronal passaging. On the day of neuronal passaging, the Matrigel droplets were aspirated and 80,000 neurons were plated per well in high density droplets (10,000 cells/µl). Following an hour to allow for cell attachment, MEA culture wells were flooded with BrainPhys culture medium (StemCell), supplemented with 1% B27, 0.5% N2, 0.02 µg/ml BDNF, 0.02 µg/ml GDNF, 0.5 mM dbcAMP, and 1 µg/ml mouse lamimin (Invitrogen). MEA neuronal cultures were fed biweekly by replacing half of the culture well’s medium volume. MEA cultures were matured for 60 days before being treated with compounds.

Spontaneous neuronal activity was measured using an Axion Biosystems Maestro Pro system. Signals were recorded using a sampling frequency of 12.5 kHz with a 3 kHz Kaiser window low-pass filter and 200 Hz high-pass filter. Action potentials were detected using the Axion Biosystems Axis Navigator software with the adaptive threshold method. Electrodes were excluded from analysis if they detected fewer than 5 spikes per minute. Quantification of neuronal firing rates was performed using the Axion Biosystems Neural Metric Tool. Neuronal activity from all activity-detecting electrodes were averaged for plotting and statistical testing. Neuronal activity was plotted in raster plots using custom scripts in Matlab (Mathworks).

### Proteomics

#### SH-SY5Y cell treatment and phosphoproteomics sample preparation

SH-SY5Y cells were plated in 6-well plates at 5×10^4^ cells/cm^2^ and grown to 80% confluence with DMEM/F12 [Gibco #11330-032] supplemented with Pen Strep [Gibco #15140-122] and 10% FBS [Cytiva #SH30071.03]. Cells were serum-starved overnight followed by treatment with 100 nM of Nt-231, Nt-237, NGF, or PBS control for 15 minutes. At 15 minutes, cells were placed on ice, washed with cold PBS twice, and removed from plate with lysis buffer (8M urea, 200 mM EPPS pH 8.5, 50 mM NaCl, Roche complete protease inhibitor tablet, Roche phosSTOP phosphatase inhibitor tablet). Cells were lysed by freeze-thaw and vortexing, clarified by centrifugation (two sequential 30-minute spins at 15,000 xG) and quantified by Pierce BCA [Thermo #23235].

Reduction was performed for each sample on 100 ug of protein from clarified lysates with 5 mM TCEP in 0.5% SDS for 10 minutes, cysteines were alkylated by addition of 10 mM iodoacetamide for 30 minutes in the dark, followed by quenching with 10 mM DTT for 10 minutes. Proteins were then isolated by SP3; in brief, 500 µg each of Mag4 and Mag6 beads [Cytiva, #65152105050250 and #45152105050250] were washed with water and added to each sample. Proteins were precipitated to beads with 80% ethanol treatment for 30 minutes. Samples were then placed on a magnetic rack and the supernatant removed. Beads were washed with 80% HPLC-grade ethanol three times and resuspended in 200 mM EPPS, pH 8.5.

Resuspended proteins were digested with 1:100 ratio of lysC:protein overnight at room temperature, followed by treatment with trypsin at 1:100 ratio of trypsin:protein at 37 °C for 6 hours. Resulting peptides were labeled with TMTpro 18-plex [Thermo, #A52045] at a ratio of 2.5:1 TMTpro:peptide for 1 hour (Li, et al. 2020). After treatment, samples were placed on a magnetic rack and supernatants from each sample were pooled, quenched with 0.5% hydroxylamine, and desalted with a 50 mg Sep-Pak [Waters, #WAT054960] (column was equilibrated with 70% ACN and 1% formic acid, sample was reconstituted in 5% ACN and 2% formic acid and applied to column, column washed with 1% formic acid, and sample was eluted with 70% ACN and 1% formic acid).

The pooled peptide sample was enriched for phosphopeptides using Fe-NTA beads [Cube, #31505-Fe] according to manufacturer’s directions. Briefly, the sample was reconstituted in 80% ACN and 0.1% TFA, applied to pre-washed Fe-NTA beads, washed three times with 80% ACN and 0.1% TFA, and eluted off the beads with 50% ACN and 2.5% NH_4_OH. Sample was immediately mixed at a 1:1 ratio with 75% ACN and 10% formic acid to reduce the pH and dried on a speedvac. The resulting enriched phosphopeptide eluate was desalted with a C18 Stage-tip prior to mass spectrometry analysis^86^.

#### Mass spectrometry data acquisition and analysis

Mass spectrometry analysis was performed on an Orbitrap Eclipse [Thermo, #FSN04-10000] equipped with an Easy-nLC [Thermo, #LC140] and a 15 cm x 75 µm C18 emitter column [Ionopticks, #ODY3-15075C18]. Peptides were separated with a 90-minute gradient from 98% buffer A (5% ACN, 0.125% formic acid) to 20% buffer B (95% ACN, 0.125% formic acid). Spectra were acquired using an SPS-MS3 method as follows: FAIMS was enabled with compensation voltages cycling between -40V, -60V, and -80V. Peptide precursors were selected from an Orbitrap MS1 scan (R: 120000, mass range: 300-2000, max inject time: 50 ms, AGC: 200%, RF lens: 30%). The top 8 precursors were isolated and fragmented for MS2 scans in the ion trap (dynamic exclusion: 90 s, isolation window: 0.5 m/z, AGC: 250%, maximum inject time: 35 ms) followed by SPS-MS3 acquisition in the Orbitrap (R: 50000, HCD NCE: 45%, max injection time: 86 ms). Sample was injected twice: once with HCD fragmentation in the MS2 ion collection (HCD NCE: 32%) and once with CID fragmentation (CID collision energy: 35%) to improve coverage^87^.

Resulting spectra were converted to mzXML using Monocle^88^ searched with Comet^89^ against the Uniprot human database with static modifications of alkylation of cysteines and TMTpro labeling to lysines and n-termini, and variable modifications of phosphorylation (STY) and oxidized methionine. Peptide spectral matches were filtered to a 1% peptide and 1% protein-level false-discovery rate. Phosphopeptide abundances were obtained based on the Orbitrap signal-to-noise quantitation of TMTpro reporter ions^90^. Quantitative data was analyzed in R: batch-correction was performed with ComBat^91^, log_2_ ratio of each treatment was calculated as log_2_(treatment/PBS control), and Pearson correlation of log_2_ ratios was calculated and reported.

#### Crystallography

Crystallization samples were prepared by concentrating to 79 mg/mL in 20 mM Tris-HCl pH 8, 150 mM NaCl. All crystallization experiments were conducted using the sitting drop vapor diffusion method. Crystallization trials were set up in200 nL drops using the 96-well plate format at 20 °C. Crystallization plates were set up using a Mosquito from SPT Labtech, then imaged using UVEX microscopes and UVEX PS-600 from JAN Scientific. Diffraction quality crystals formed in 0.2 M Magnesium chloride hexahydrate, 0.1 M Tris pH 8.5 and 25% w/v Polyethylene glycol 3,350 for 231_C2_TrkA; in 28 % w/v acrylic acid/maleic acid copolymer (50:50), sodium salt and 0.1 M HEPES-NaOH pH 6.5 for C2-01018; and in 15% (w/v) PEG 1500 for C2-02802.

Diffraction data was collected at the Advanced Photon Source (APS) on 24ID-C for C2-01018 and C2-02802. For 231_C2_TrkA (Nt-231) diffraction data collected on NSLS2 FMX. X-ray intensities and data reduction were evaluated and integrated using XDS^69^ and merged/scaled using Pointless/Aimless in the CCP4 program suite^69,70^. Starting phases were obtained by molecular replacement using Phaser^71^ using the designed model for the structures. Following molecular replacement, the models were improved using phenix.autobuild^72^; efforts were made to reduce model bias by using simulated annealing. Structures were refined in Phenix^72^. Model building was performed using COOT^73^. The final model was evaluated using MolProbity^74^. Data collection and refinement statistics are recorded in **Table S1**. Data deposition, atomic coordinates, and structure factors reported in this paper have been deposited in the Protein Data Bank (PDB), http://www.rcsb.org/, with accession codes **9DT9**, **9DTD**, and **9DTE.**

## Quantification and statistical analysis

Details of the quantification and statistical analysis can be found in the Figure legend and method details section. Unless otherwise noted, all data was plotted and fitted using Origin Pro 10 or GraphPad Prism 10. Surface Plasmon Resonance data was analyzed via the Biacore Insight Evaluation Software and exported for further processing with Origin Pro 10. Mass Photometry data was plotted and fitted using a Gaussian fit with the DiscoverMP software. Analysis of neurite outgrowth in neuroblastoma cell lines was performed with the Neurotrack Analysis Software Module of the IncuCyte Live-Cell Analysis System. Differential expression analysis for RNASeq Data was performed using R/Bioconductor package DESeq2 and edgeR. Heatmaps were generated with the R package pheatmap or the Python library matplotlib. Quantification of neuronal firing rates in the MEA assay was performed using the Axion Biosystems Neural Metric Tool. Starting phases were generated by molecular replacement using Phaser with the design model as input structure. Models were improved using phenix.autobuild and further refined in Phenix. Model building was performed using COOT and evaluated using MolProbity.

## Supplementary Figures

**Supplementary Figure 1:**
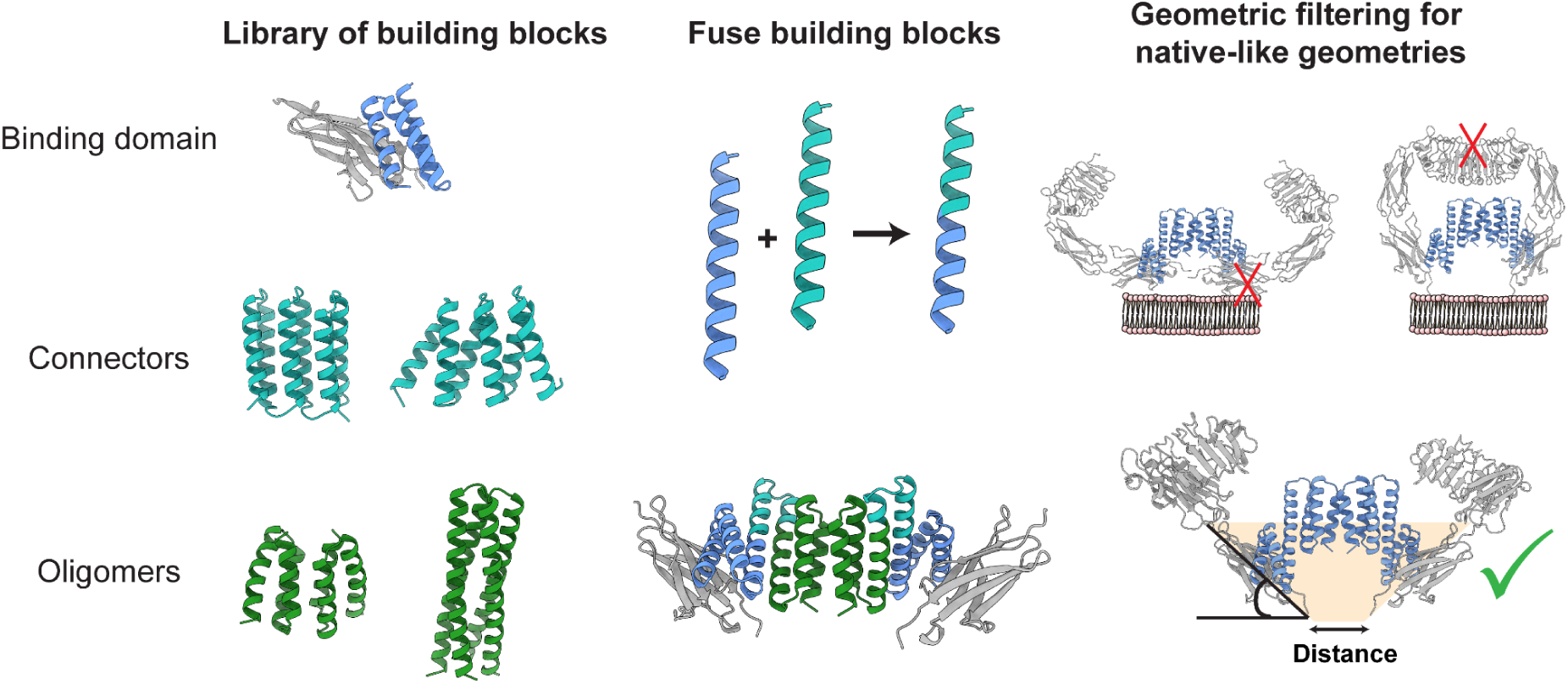
Design Protocol, designing for native like geometries. The first step in the protocol selects building blocks from a library (TrkA binding domain, Helical Repeat Connectors/DHRs, C2-Oligomers). Those building blocks are then fused along secondary structure elements. The last step of the backbone generation is to filter for native-like outputs considering geometric parameters, such as native-like angles of TrkA domain 3 (20^∘^ < θ < 80^∘^) towards the membrane and clash checks between the two receptors. Backbone design is then followed by Rosetta or MPNN-based sequence design.

**Supplementary Figure 2:**
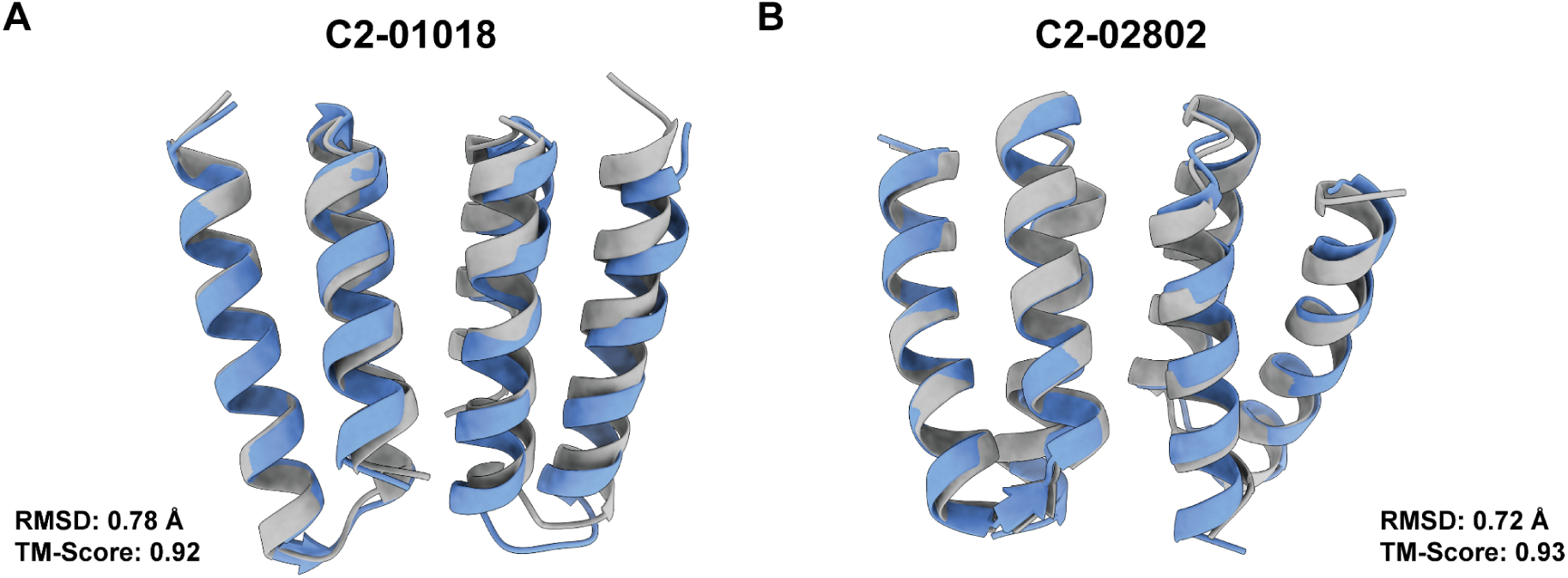
Designed C2 Oligomers match crystal structure. (A) The crystal structure of C2-01018 (gray) matches the design model (blue) with subangstrom accuracy (PDB ID: 9DTD). (B) The crystal structure of C2-02802 (gray) matches the design model (blue) (PDB ID: 9DTE).

**Supplementary Figure 3:**
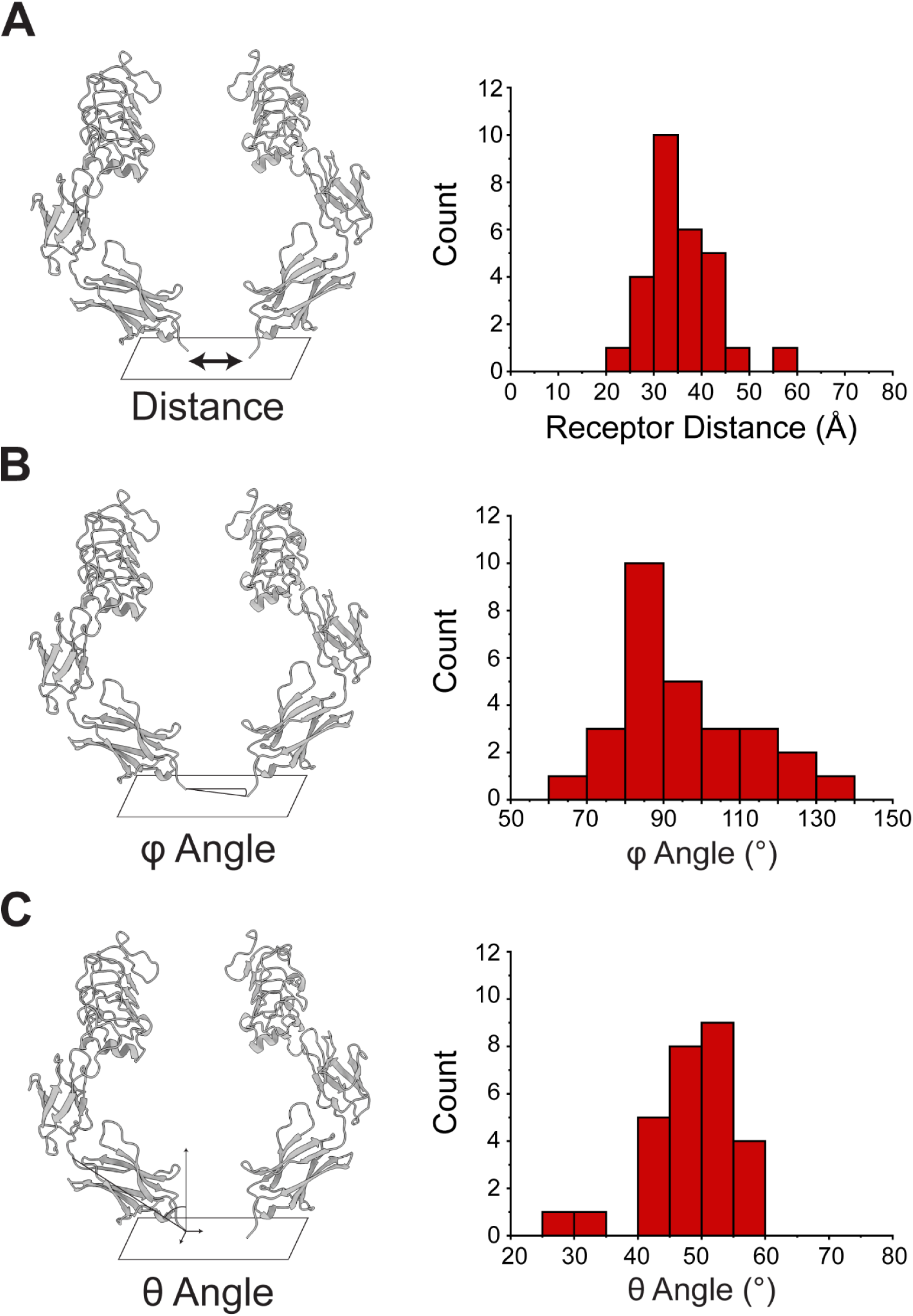
Diversity of designs templating the receptor architecture. (A) receptor distance (B) φ angle (C) θ angle.

**Supplementary Figure 4:**
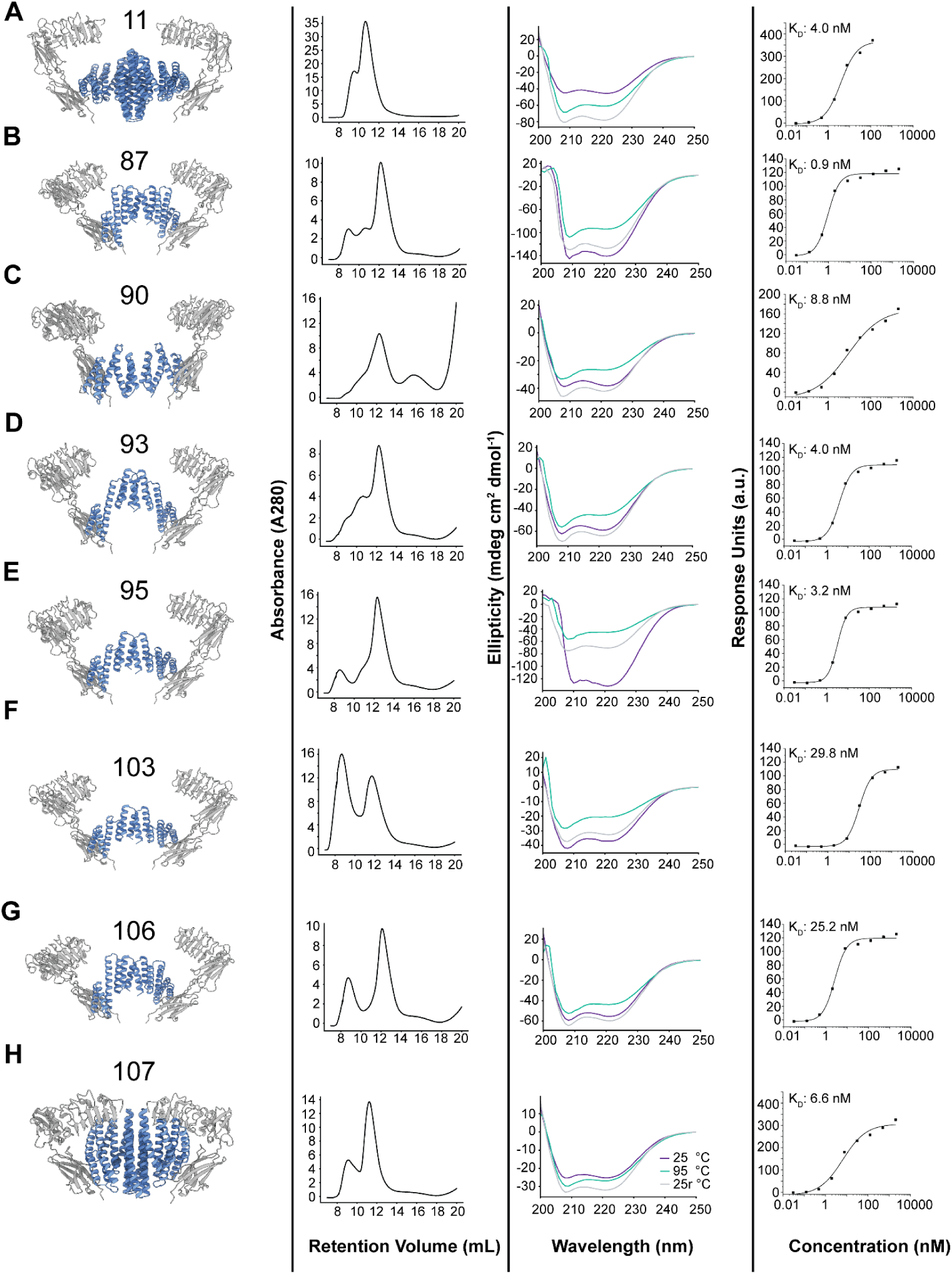
Designed cyclic oligomers that engage the TrkA receptor. 1st column: cartoon model, 2nd column: size-exclusion purification profile, 3rd column: CD spectra including different temperatures, 4th column: Dose-response curve of target binding on SPR.

**Supplementary Figure 5:**
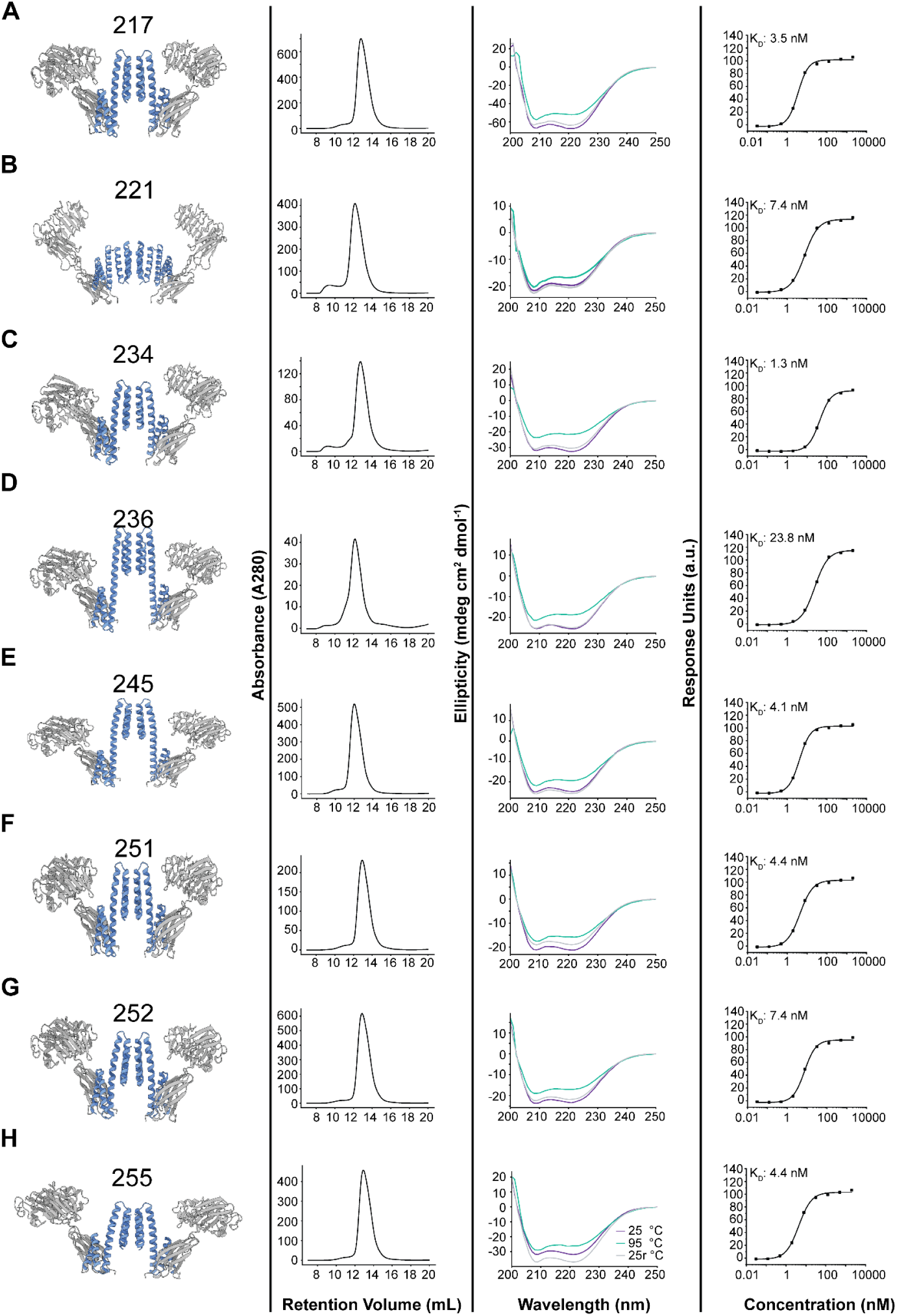
Deep-Learning designed oligomers. 1st column: cartoon model, 2nd column: size-exclusion purification profile, 3rd column: CD spectra including different temperatures, 4th column: Dose-response curve of target binding on SPR.

**Supplementary Figure 6:**
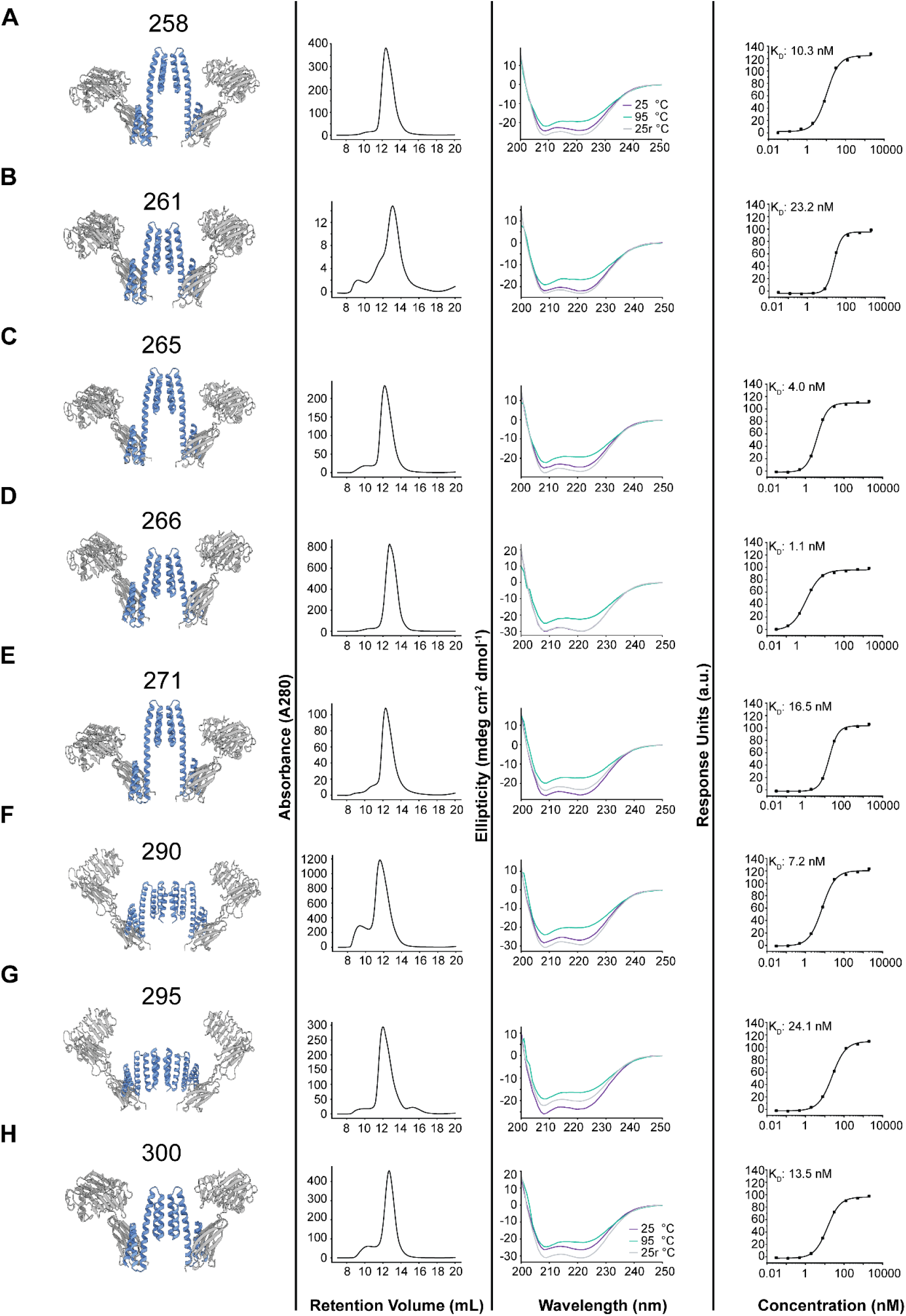
Deep-Learning designed oligomers. 1st column: cartoon model, 2nd column: size-exclusion purification profile, 3rd column: CD spectra including different temperatures, 4th column: Dose-response curve of target binding on SPR.

**Supplementary Figure 7:**
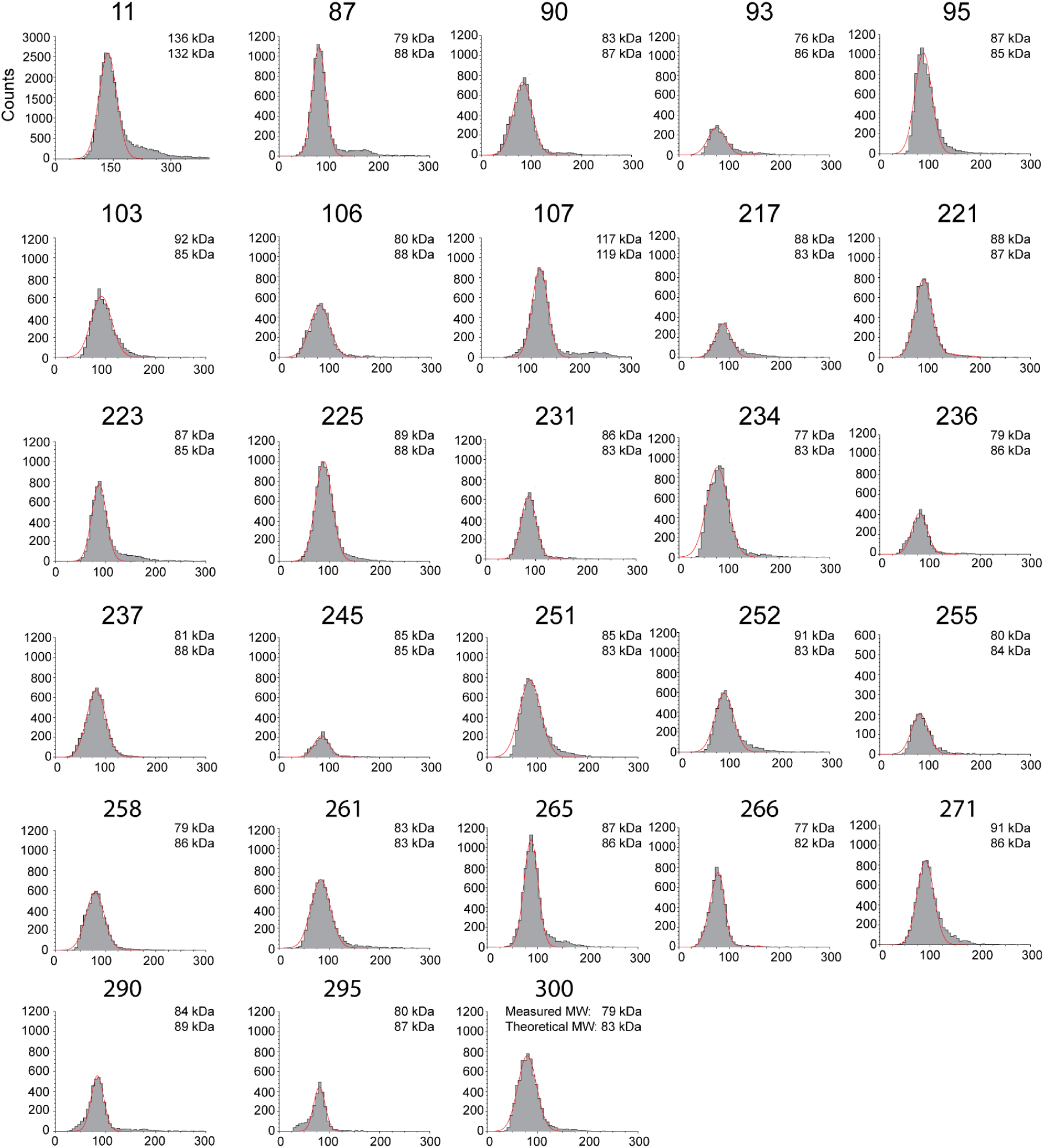
Mass-Photometry analysis of C2-oligomeric constructs with GFP-mass tag.

**Supplementary Figure 8:**
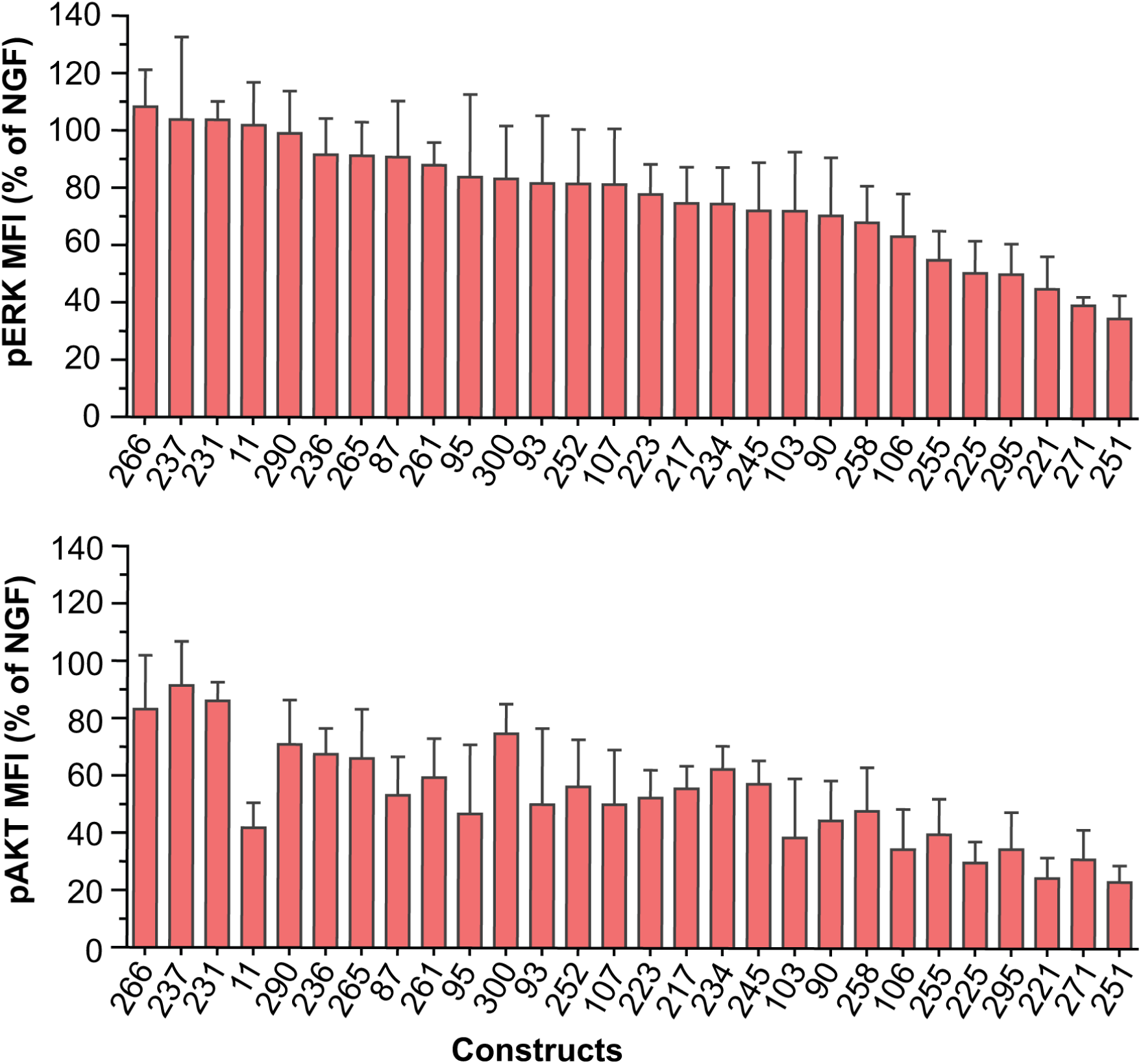
E_max_ values for pERK (top) and pAKT (bottom) activation for the individual constructs in TF-1 cells. Error bars represent SD from three independent biological repeats.

**Supplementary Figure 9:**
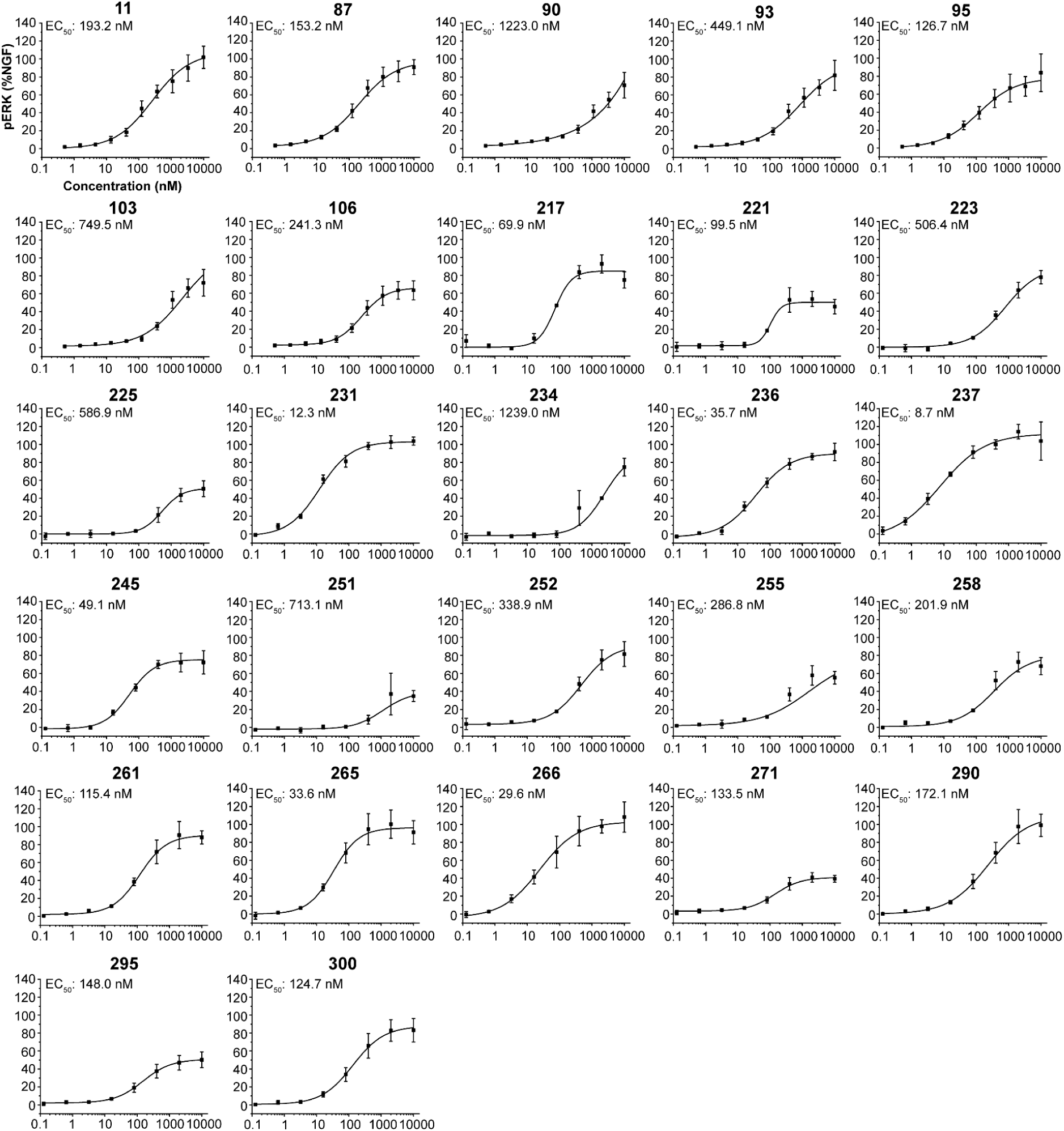
Dose-response curve for ERK signaling for all measured constructs in TF-1 cells. Individual plots of data from Figure 3. Error bars represent SD from three independent biological repeats.

**Supplementary Figure 10:**
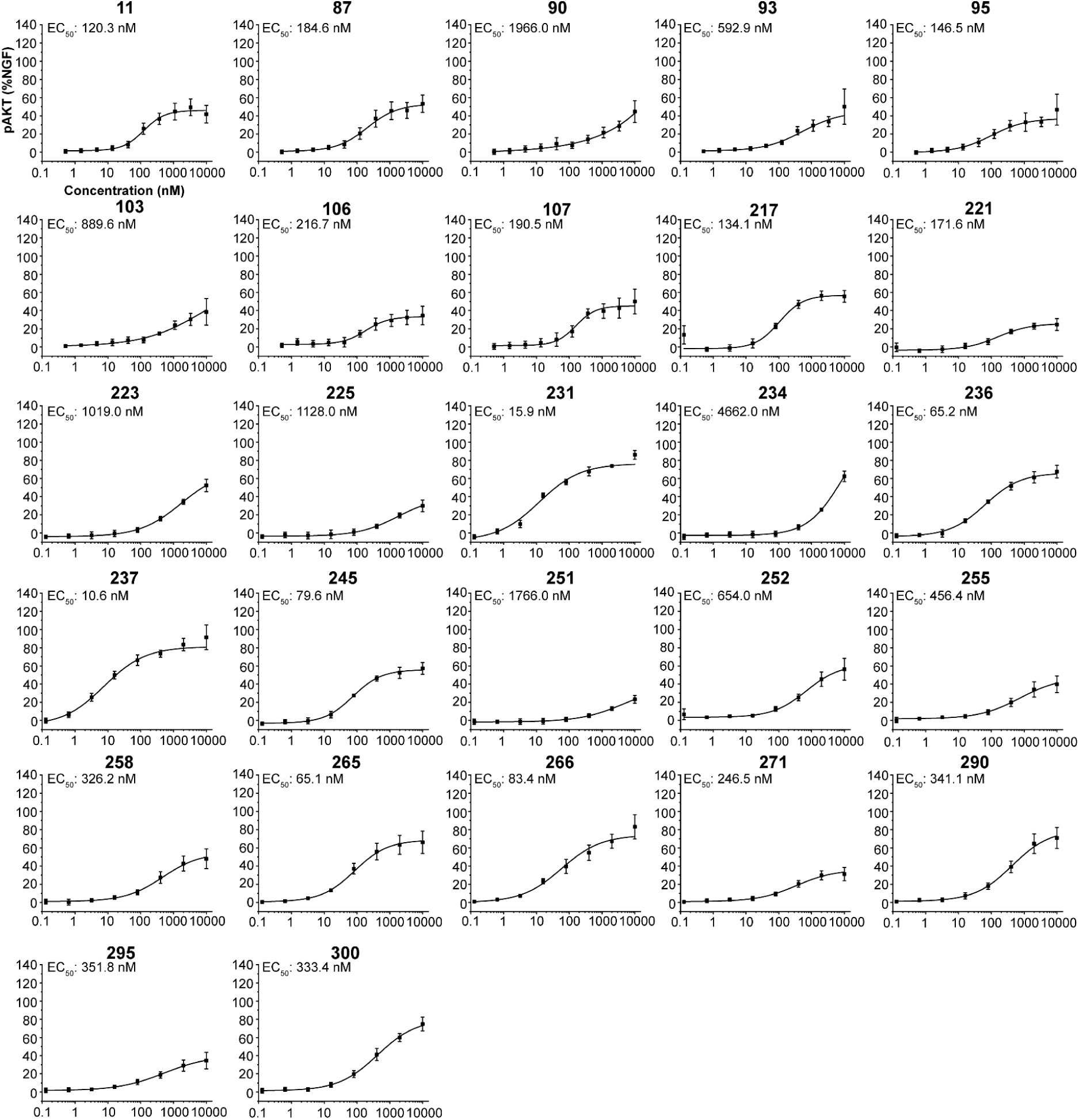
Dose-response curve for AKT signaling for all measured constructs in TF-1 cells. Individual plots of data from Figure 3. Error bars represent SD from three independent biological repeats.

**Supplementary Figure 11:**
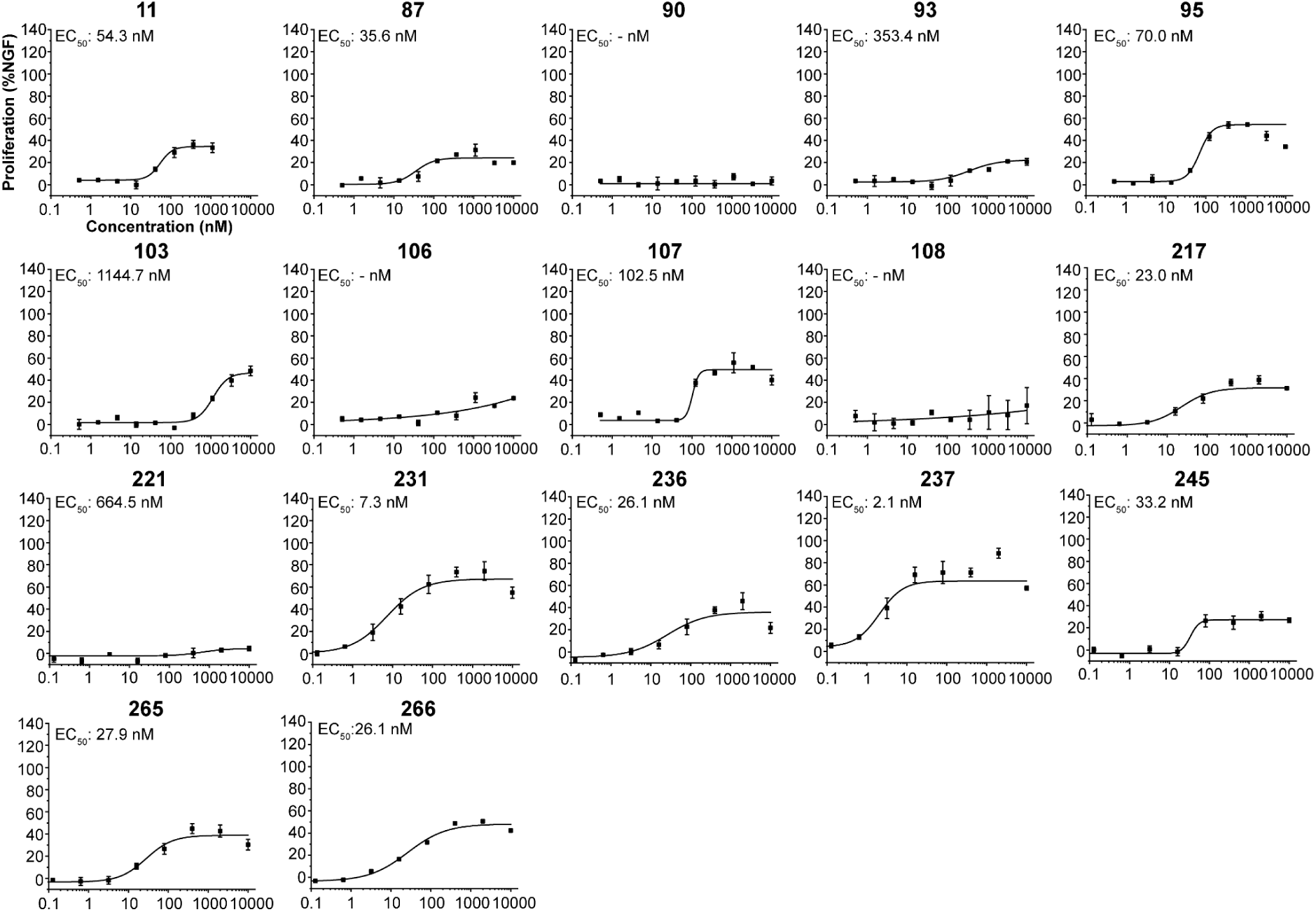
Dose-response curve for Proliferation signaling for all measured constructs in TF-1 cells. Error bars represent SD from three independent biological repeats.

**Supplementary Figure 12:**
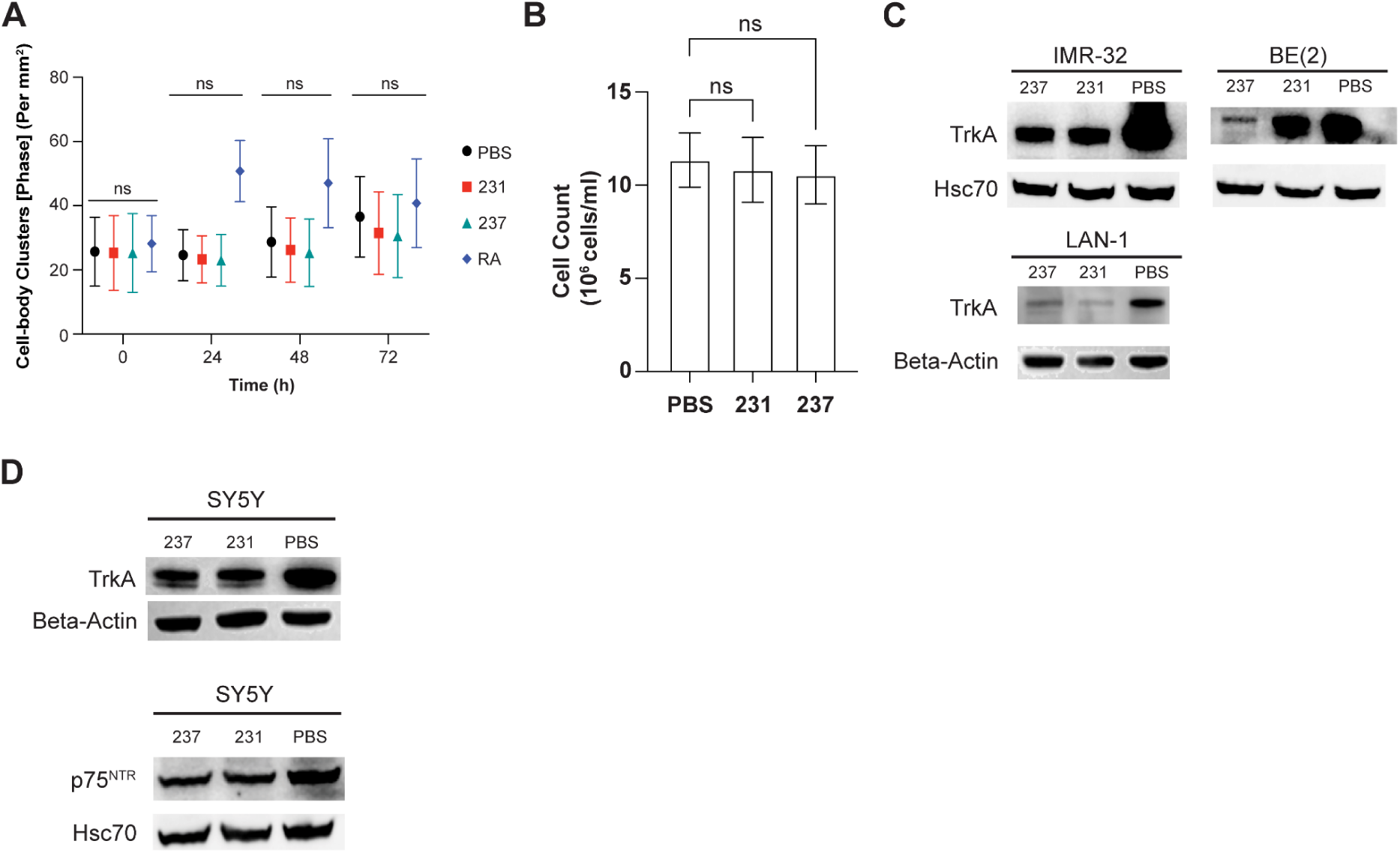
Neuroblastoma differentiation. (A) Comparison of detected cell-body clusters of SH-SY5Y cells treated with PBS, Nt-231, Nt-237 or RA. (B) Proliferation of SH-SY5Y cells after stimulation with constructs. (C) Downregulation of TrkA receptor in 3 different neuroblastoma cell lines. (D) Expression levels of TrkA and p75^NTR^ in SH-SY5Y cells. Error bars represent SD from three independent biological repeats.

**Supplementary Figure 13:**
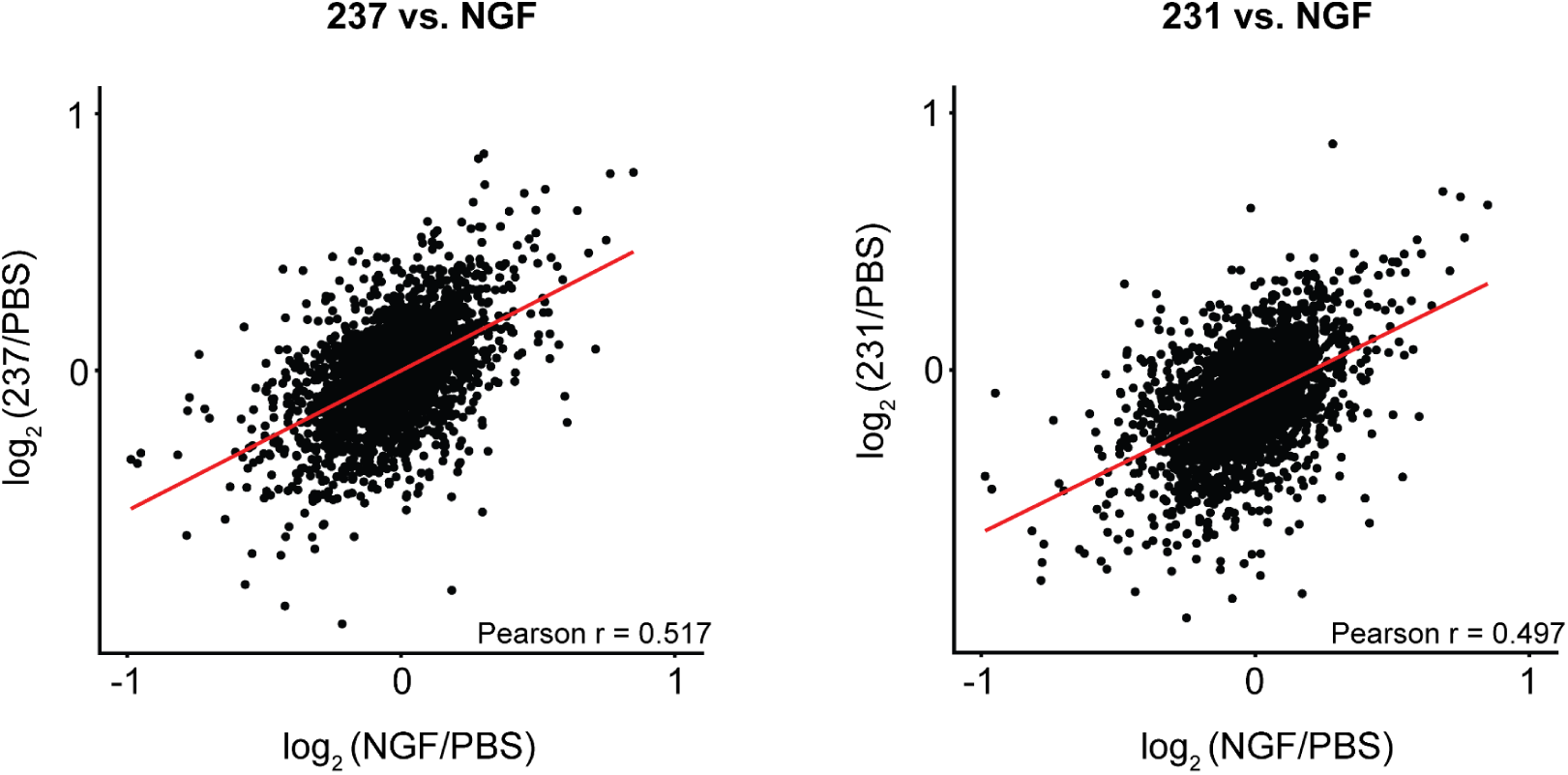
Phospho-Proteomics of treated SH-SY5Y cells. Phospho-Proteomics shows moderate correlation of the two neotrophins to NGF signaling in SH-SY5Y cells after 15 min of treatment with 100 nM.

**Supplementary Figure 14:**
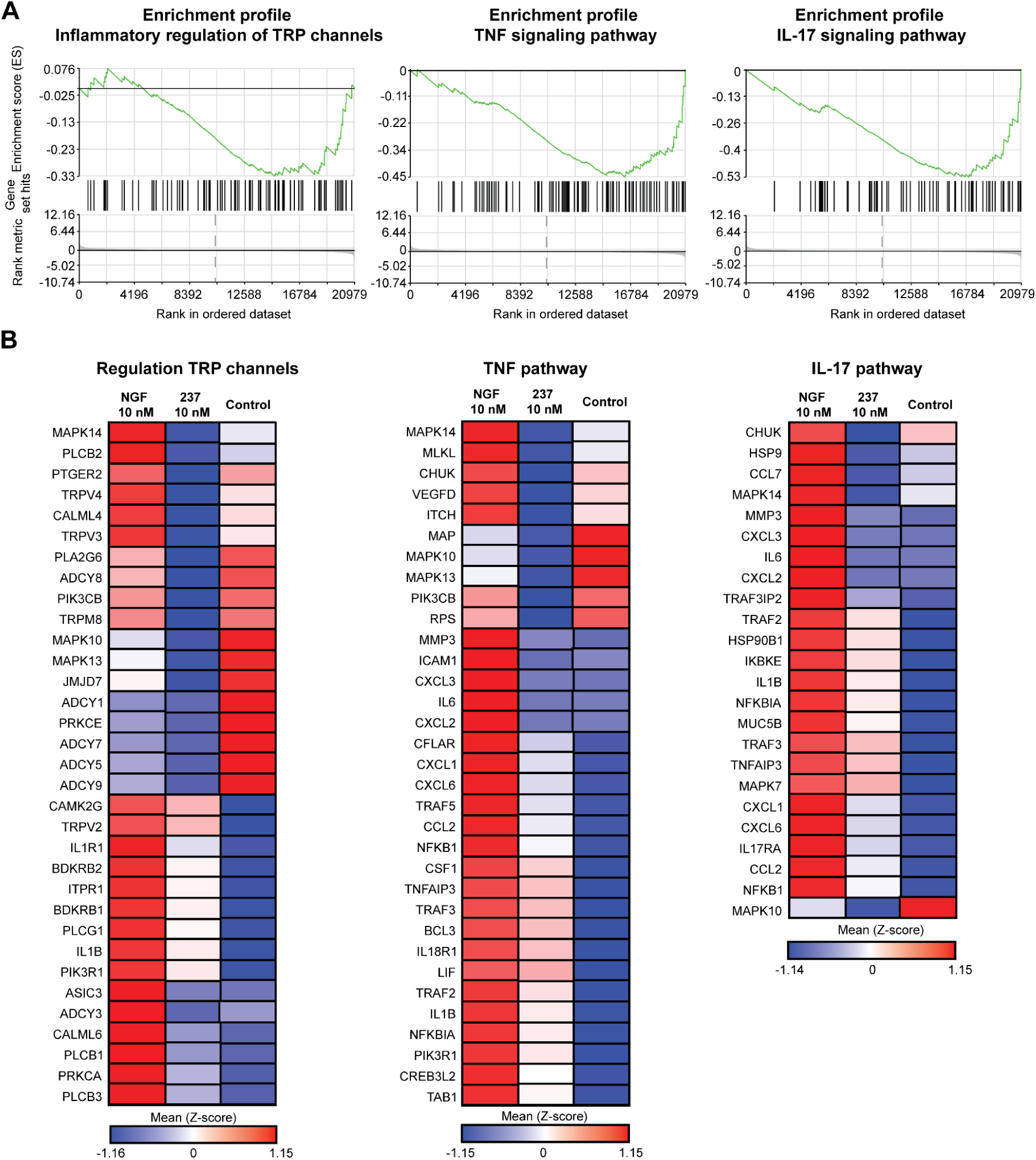
mRNA sequencing analysis of TrkA and p75^NTR^ expressing neurons. (A) Enrichment profiles between 237 and NGF treated iNs. (B) Heatmaps of gene expression difference for leading edge genes of selected pathways.

## Supplementary Tables

**Supplementary Table 1.** Data collection and refinement statistics

|  | **231_C2_TrkA** (PDB Code: 9DT9) | **C2-01018** (PDB Code: 9DTD) | **C2-02802** (PDB Code: 9DTE) |
| --- | --- | --- | --- |
| **Data collection** |  |  |  |
| Space group | $P 2_1$ | $P 3_2 2 1$ | $R 3 :H$ |
| Cell dimensions |  |  |  |
| $a, b, c$ (Å) | 41.47, 60.46 51.88 | 69.31, 69.31, 49.41 | 56.43, 56.43, 96.62 |
| $\alpha, \beta, \gamma$ (°) | 90, 94.09, 90 | 90, 90, 120 | 90, 90, 120 |
| Resolution (Å) | 51.75 - 2.2 (2.29 - 2.2) | 60.02 - 3.10 (3.21 - 3.10) | 43.61 - 1.49 (1.54 -1.49) |
| $R_{\text{merge}}$ | 0.117 (0.644) | 0.2874 (0.7225) | 0.067 (1.185) |
| $I / \sigma I$ | 8.69 (2.10) | 3.58 (1.37) | 11.88 (0.87) |
| CC1/2 | 0.997 (0.851) | 0.968 (0.844) | 0.997 (0.469) |
| Completeness (%) | 99.36 (99.66) | 97.48 (96.94) | 99.71 (99.14) |
| Redundancy | 4.7 (4.7) | 5.0 (5.2) | 5.1 (4.8) |
| **Refinement** |  |  |  |
| Resolution (Å) | 51.75 - 2.2 (2.29 - 2.2) | 60.02 - 3.10 (3.21 - 3.10) | 43.61 - 1.49 (1.54 -1.49) |
| No. reflections | 13024 (1449) | 4715 (2618) | 18703 (1843) |
| $R_{\text{work}} / R_{\text{free}}$ | 0.2259 (0.2693)/ 0.2866 (0.3202) | 0.2547 (0.2701)/ 0.3159 (0.4064) | 0.1915 (0.3241)/ 0.2117 (0.3305) |
| No. atoms |  |  |  |
| Protein | 1816 | 954 | 984 |
| Water | 109 | n/a | 46 |
| B-factors |  |  |  |
| Protein | 43 | 109 | 43 |
| Water | 42 | n/a | 62 |
| R.m.s. deviations |  |  |  |
| Bond lengths (Å) | 0.002 | 0.004 | 0.002 |
| Bond angles (°) | 0.330 | 0.573 | 0.370 |

\*A single crystal was used for each structure in the data collection. Values in parentheses are for the highest-resolution shell.

**Supplementary Table 2:** Signaling Data of constructs.

| Construct | ERK |  | AKT |  | Affinity |
| --- | --- | --- | --- | --- | --- |
|  | EC<sub>50</sub> (nM) | E<sub>max</sub> (%NGF) | EC<sub>50</sub> (nM) | E<sub>max</sub> (%NGF) | K<sub>D</sub> (nM) |
| Nt-11 | 193.2 | 96.7 | 120.3 | 47.6 | 4.0 |
| Nt-87 | 153.2 | 91.2 | 184.6 | 52.1 | 0.9 |
| Nt-90 | 1223.0 | 76.6 | 1966.0 | 50.0 | 8.8 |
| Nt-93 | 449.1 | 81.3 | 592.9 | 46.9 | 4.0 |
| Nt-95 | 126.7 | 76.6 | 146.5 | 40.3 | 3.2 |
| Nt-103 | 749.5 | 79.8 | 889.6 | 40.2 | 29.8 |
| Nt-106 | 241.3 | 67.3 | 216.7 | 35.0 | 25.2 |
| Nt-107 | 267.7 | 81.7 | 190.5 | 48.4 | 6.6 |
| Nt-217 | 69.9 | 88.4 | 134.1 | 58.7 | 3.5 |
| Nt-221 | 99.5 | 53.6 | 171.6 | 25.1 | 7.4 |
| Nt-223 | 506.4 | 81.2 | 1019.0 | 56.8 | 18.8 |
| Nt-225 | 586.9 | 54.7 | 1128.0 | 33.3 | 1.7 |
| Nt-231 | 12.3 | 101.5 | 15.9 | 76.0 | 12.7 |
| Nt-234 | 1239.0 | 80.2 | 4662.0 | 92.5 | 1.3 |
| Nt-236 | 35.7 | 88.4 | 65.2 | 64.5 | 23.8 |
| Nt-237 | 8.7 | 105.4 | 10.6 | 82.1 | 8.2 |
| Nt-245 | 49.1 | 74.7 | 79.6 | 56.2 | 4.1 |
| Nt-251 | 713.1 | 41.5 | 1766.0 | 27.2 | 4.4 |
| Nt-252 | 338.9 | 85.5 | 654.0 | 59.2 | 7.4 |
| Nt-255 | 286.8 | 60.5 | 456.4 | 41.4 | 4.4 |
| Nt-258 | 201.9 | 74.5 | 326.2 | 49.4 | 10.3 |
| Nt-261 | 115.4 | 92.0 | 207.2 | 59.9 | 23.2 |
| Nt-265 | 33.6 | 98.3 | 65.1 | 65.7 | 4.0 |
| Nt-266 | 29.6 | 101.7 | 83.4 | 74.5 | 1.1 |
| Nt-271 | 133.5 | 43.0 | 246.5 | 32.4 | 16.5 |
| Nt-290 | 172.1 | 102.0 | 341.1 | 73.7 | 7.2 |
| Nt-295 | 148.0 | 50.7 | 351.8 | 34.8 | 24.1 |
| Nt-300 | 124.7 | 86.0 | 333.4 | 73.9 | 13.5 |

**Supplementary Table 3:**
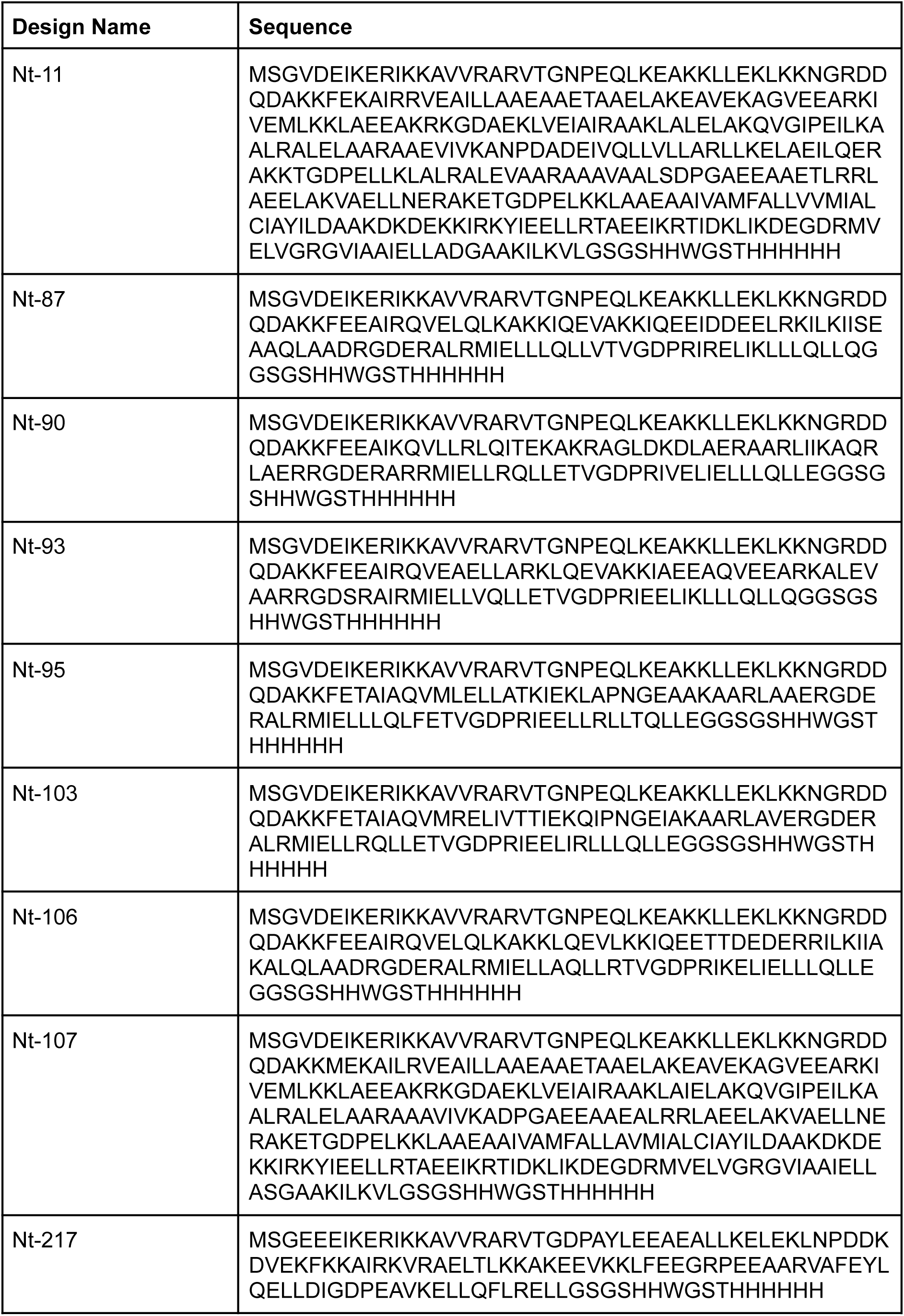
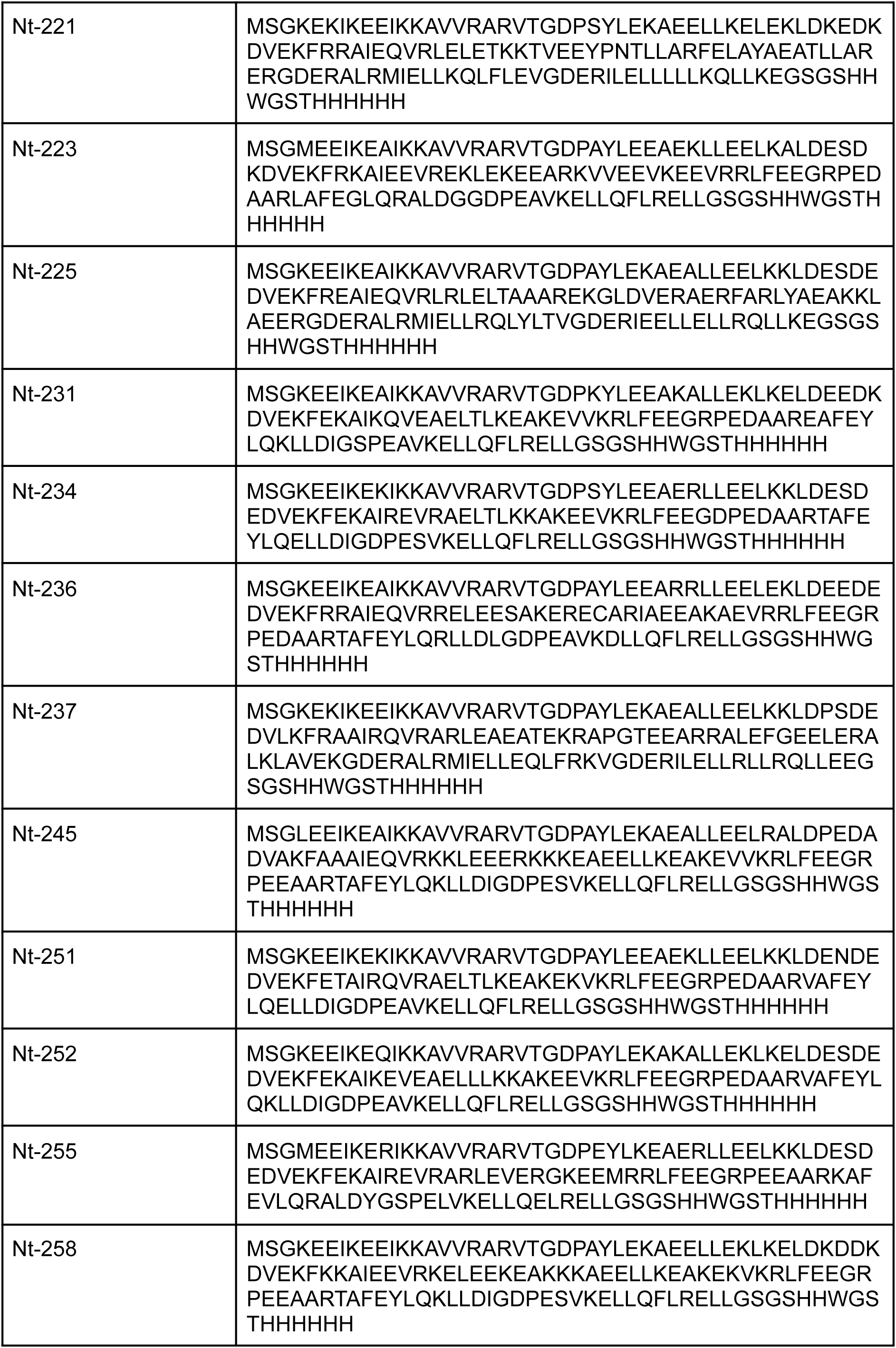
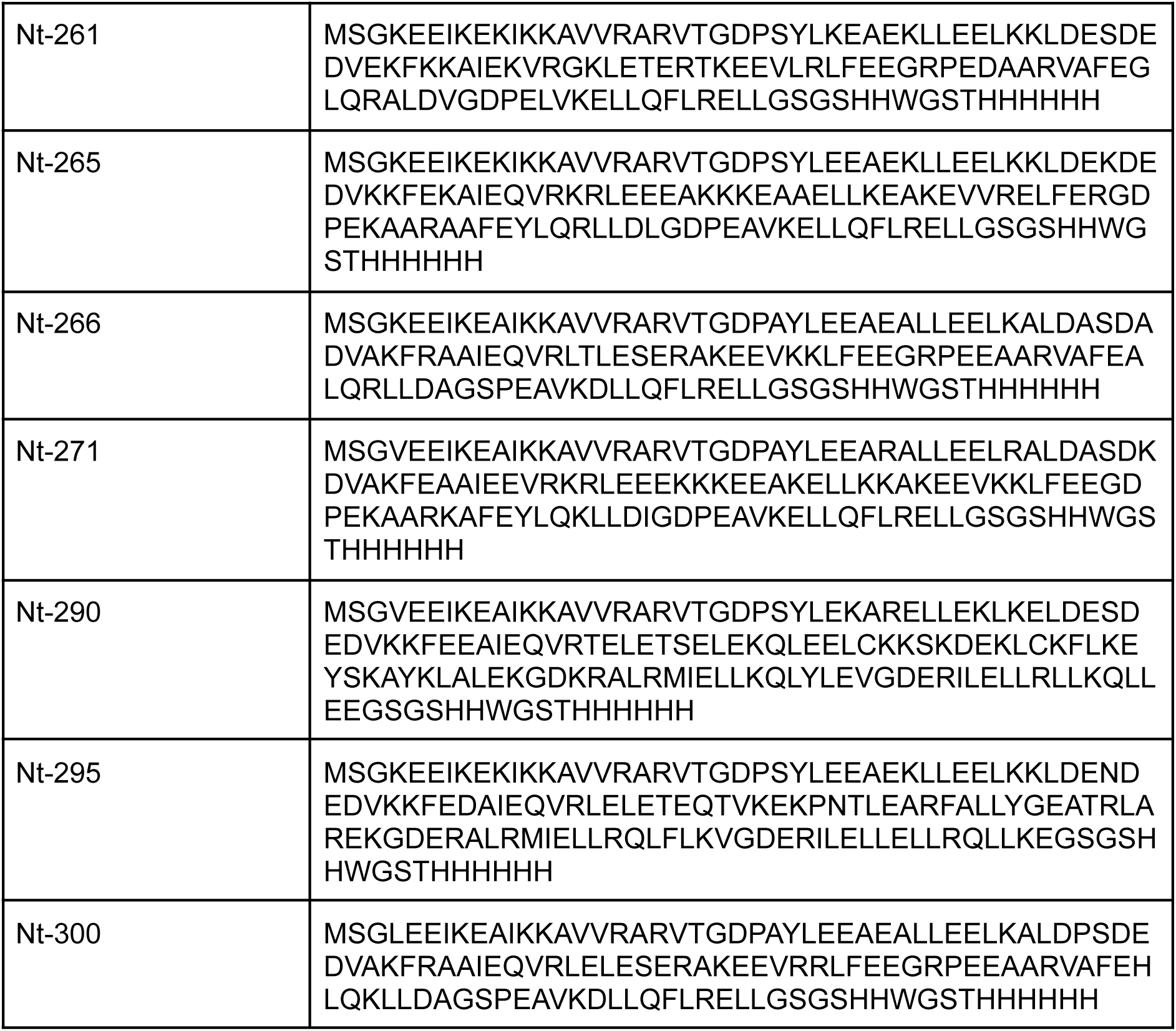
Sequences of designed proteins

**Supplementary Table 4:**
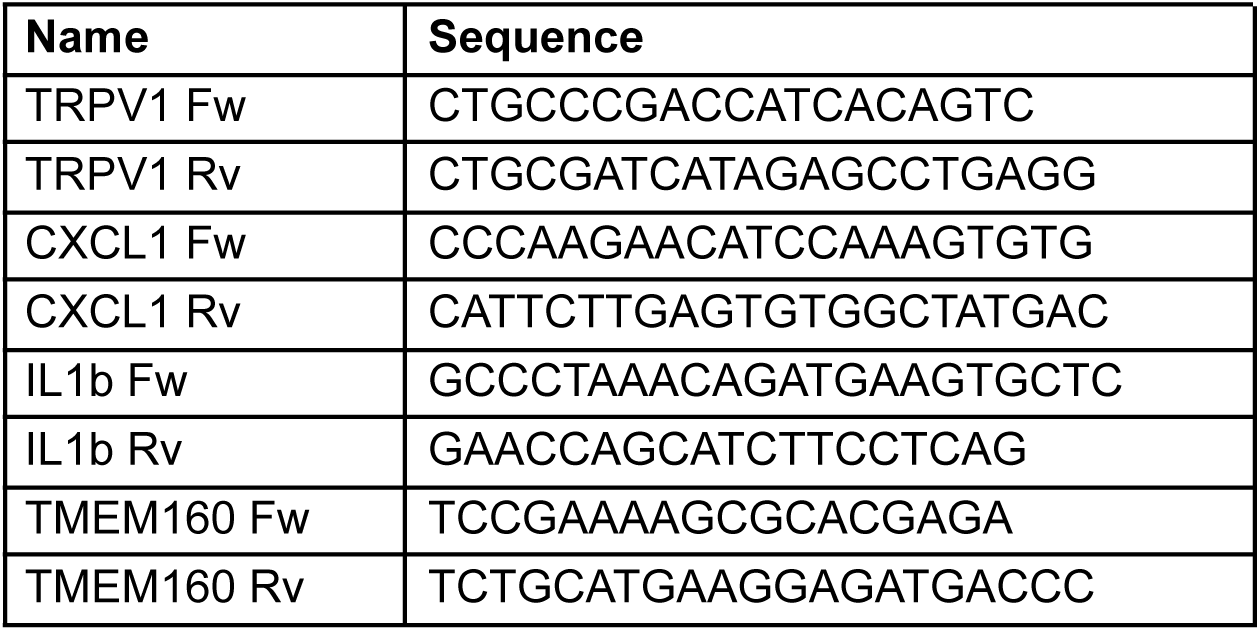
qPCR Primers

